# Temporal Contrast Enhancement in Auditory and Nociceptive Processing

**DOI:** 10.64898/2025.12.16.694610

**Authors:** Jakob Poehlmann, Tibor M. Szikszay, Luisa Luebke, Waclaw M. Adamczyk, Kerstin Luedtke, Malte Wöstmann

**Affiliations:** Institute of Health Sciences, Department of Physiotherapy, Pain and Exercise Research Luebeck (P.E.R.L.), University of Luebeck, Lübeck, Germany; Center of Brain, Behavior and Metabolism (CBBM), University of Luebeck, Lübeck, Germany; Laboratory of Pain Research, Institute of Physiotherapy and Health Sciences, Academy of Physical Education, Katowice, Poland; Department of Psychology, University of Luebeck, Lübeck, Germany

**Keywords:** Temporal Filtering, Temporal Contrast Enhancement, Offset Analgesia, Auditory System, Pain, Sensory Processing

## Abstract

Temporal contrast enhancement (TCE), also termed offset analgesia, describes a temporal filtering mechanism whereby a small decrease in stimulus intensity produces a disproportionately large reduction in perceived pain. Although TCE is considered a robust marker of endogenous pain modulation, its underlying mechanisms remain unresolved. It is unclear whether TCE reflects a nociceptive-specific modulatory process or a supramodal temporal filtering mechanism that generalizes to non-painful but aversive sensory stimulation. In this study, healthy and pain-free participants were enrolled in two experiments: a behavioral study (n=33) or a neurophysiological study (n=29). Continuous pain-ratings, electroencephalography (EEG) and pupillometry data were collected. A conventional TCE paradigm was applied using either noxious heat delivered via a thermal contact stimulator or unpleasant auditory stimulation delivered through over-ear headphones. Both noxious heat and unpleasant sounds induced behavioral TCE-effects (p<0.01), indicating a supramodal mechanism with modality-specific temporal dynamics, which was further supported by the absence of a significant correlation of TCE-effects across modalities (p>0.05). In the second experiment, noxious heat – but not unpleasant sound – resulted in decreased power of neural alpha oscillations (∼10 Hz, p<0.05) and increased pupil size (p<0.05) potentially indicating bottom-up modulation of the autonomic nervous system and a release of neural inhibition, respectively. However, subjectively experienced TCE-effects were not reflected in neurophysiological correlates. These findings support the view of TCE as a supramodal temporal filtering mechanism characterized by modality-specific temporal dynamics, for which no corresponding signatures are found in EEG or pupillometry.

## 1. Introduction

Human environments contain a plethora of sensory inputs. Sensory integration is a complex process dependent on stimulus intensity, salience and context [85]. Therefore, diverse filtering mechanisms covering spatiotemporal aspects are required to selectively enhance specific features of stimuli, evaluate their relevance, and modulate perceptual processing.[12,54]. In the context of auditory perception, temporal summation - alternatively referred to as temporal integration [56] or temporal sound summation [6] - describes the perceived increases in loudness when a tone of constant volume is presented for longer durations. Temporal filtering of auditory percepts is of utmost importance, due to its relevance in vocalization and speech recognition [43], as well as in avoiding uncomfortably loud sounds. Temporal summation likewise characterizes nociceptive processing [11], where adaptive temporal filtering is vital for averting tissue injury and protecting the organism. Evidence suggests that temporal filtering processes share similar mechanisms across modalities and sensory systems [7,16,23,61]. However, the extent to which temporal filtering in auditory and nociceptive processing is based on common mechanisms remains unclear.

Temporal contrast enhancement (TCE), a putative component of the endogenous pain modulatory system, is conceptualized as a temporal filtering process whose mechanisms remain unclear [51,81,84]. Commonly referred to as offset analgesia, it is characterized by a disproportionally large reduction in perceived pain following a slight decrease in noxious stimulus intensity [22,81,82]. A variety of mechanisms have been postulated as potential mediators of TCE. Several brain imaging studies using functional magnetic resonance imaging (fMRI) showed altered activity in cortical [15,51,81] and subcortical structures [5,51,63,81,84].

Interestingly, studies investigating the temporal dynamics of TCE using electroencephalography (EEG) are scarce. Research has focused on EEG during applications of tonic or phasic pain, investigating alpha-band oscillations (∼10Hz), which consistently showed reduced alpha power during pain [52,53,58]. This might reflect functional inhibition of situational irrelevant information [14,29,30,32,64], which is already well-established in auditory attention research [62,77,78]. Furthermore, alpha oscillatory power has been shown to relate to temporal expectations [60,76,80] and stimulus predictability [79], making it a suitable neural marker for examining how expectations of increasing pain or sound intensity shape temporal filtering. Additional objective measures have been used to characterize neurophysiological correlates of pain and auditory processing. Changes in pupil diameter are closely related to autonomic nervous system activity and have been linked to perceived stimulus intensity and arousal, particularly in response to painful stimuli [10,34]. In auditory research, pupil dilation has been associated with loudness of the stimuli, salience, and aversiveness [37,74,83], highlighting its relevance for tracking perceptual and attentional processes.

Here, it is hypothesized that the effects of TCE on noxious stimuli may not reflect pain-specific processes but rather could be indicative of a general temporal filtering mechanism in response to salient stimuli. To determine whether observed effects in pain are modality-specific or merely reflect general arousal, prior studies have employed salience-matched control stimuli; nevertheless the evidence still remains equivocal [27,35,48,58,72]. Therefore, we conducted a multimodal study to determine whether TCE is specific to nociception, using painful heat and aversive auditory stimulation within a standard TCE paradigm.

## 2. Materials and methods

Two experiments were conducted: the first investigated behavioral measures of TCE, while the second additionally incorporated (neuro-)physiological measures. Behavioral outcomes were used in confirmatory analyses, whereas (neuro-)physiological responses derived from EEG and pupillometry were examined exploratorily to support understanding the mechanistic basis of temporal filtering mechanisms.

### 2.1. Experiment 1: Behavioral investigation of the TCE effect in auditory stimulation

The aim of the first experiment was to investigate if it is possible to elicit TCE behaviorally using auditory stimulation. The study was approved by the Ethics Committee of the University of Luebeck (2023-633) and conducted in accordance with the Declaration of Helsinki. The methodology was preregistered on the Open Science Framework (OSF; https://osf.io/643kg).

#### 2.1.1. Participants

Participants (n = 33, 22 females; age = 23.7, Standard deviation (SD) = 6.0) were included if they subjectively reported being healthy and pain-free on the day of the procedure. Exclusion criteria were chronic pain (> 3 months) within the last two years, diagnosed systemic, neurological, cardiovascular or psychiatric diseases and being with diagnosed hearing loss or tinnitus. All participants were asked not to take any painkillers, consume alcohol or undertake any strenuous physical activity 24 hours before participating in the study. To characterize the study sample age, sex at birth, body mass (kg), stature (cm), handedness and general fear of heat pain and fear of loud noises on a numerical rating scale from 0 (no fear) to 100 (highest possible fear) were recorded for each participant. To measure noise sensitivity the Weinstein Noise Sensitivity Scale (WNSS) was used [86]. In addition, the Pain Vigilance and Awareness Questionnaire (PVAQ) was used to measure attention to pain and assesses awareness, consciousness, vigilance, and observation of pain [44] and the Pain Catastrophizing Scale (PCS) was used to assess the catastrophizing behavior [45].

#### 2.1.2. Equipment

All heat stimuli were applied with a thermal contact stimulator (TCS; André Dufour, University of Strasbourg, France), a CE certified, validated stimulation device developed by QST.lab [71]. The probe of the TCS weighs 440g and has a temperature range of 0°C to 60°C, adjustable at 0.1°C intervals. The maximum temperature rise and fall rate is 100°C/second. It has a total stimulation zone of 9 cm^2^ (five equal stimulation zones, each 0.74 x 2.4cm = 8.88cm^2^) and was applied to the left forearm. For auditory stimulation, a 1000-Hz sine wave tone was generated at a sampling rate of 44.1 kHz using Adobe Audition (Adobe Systems Software, Dublin, Republic of Ireland). The sound was presented using over-ear headphones (PXC 550-II, Sennheiser, Wedemark, Germany). Probing pain intensity was conducted with a Python-based eVAS [20] and was adapted using a scale from Bushnell et al. [47]. The eVAS was displayed on a computer screen and consisted of a left anchor (0/200), middle anchor (100/200), and right anchor (200/200). For heat stimulation, the left anchor represented “no sensation”, meaning participants were instructed to keep the eVAS slider in this initial position when not feeling any sensation of warmth or pain. As soon as any warmth sensation occurred, participants indicated this by moving the cursor to the right. When the heat sensation was perceived as being painful, they had to cross the “heat pain threshold” (100) set in the middle of the eVAS. The right portion of the eVAS (100–200) represented the range from painful to “worst pain imaginable” (200). For sound stimulation, the eVAS was constructed similarly, with the left anchor (0) representing “no sound” perception. Participants moved the cursor with increasing or decreasing perceived sound intensity. When the sound was perceived as being uncomfortable, they crossed the middle anchor representing the “discomfort threshold” (100). The right anchor represented “maximum discomfort imaginable” (200).

#### 2.1.3. Familiarization and calibration procedure

Before attending the main part of the experiment, participants were familiarized to the rating procedure and stimulus intensities. For this, they received three different stimulus intensities, a high (48°C, 100dB), low (33°C, 49dB) and an intermediate (40°C, 82dB) level of intensity for thermal and auditory stimulation, respectively. Afterwards, the stimulation intensities for each modality were calibrated, using a staircase procedure (S1 Fig. in the Supporting Information). Participants were asked to continuously rate their sensory experience using the eVAS scale. The calibration consisted of ascending stimulation intensities, where each stimulation was applied for ten seconds, followed by ten seconds of either no stimulation (0dB) or baseline temperature (32°C) depending on the applied modality. The stimulation started at either 33°C or 49dB and increased in 1°C or 3dB steps up until the highest stimulation intensity of 48°C or 100dB was reached. This procedure was repeated for each modality with the modalities being alternated. The stimulation intensities were then derived from the second iteration of each modality by using stimulation intensities that produced perceived pain intensities of 150/200 and 175/200 on the eVAS. These values were used for the initial stimulation intensity (T1) and increase in stimulation intensity (T2) in offset trials (see Experimental paradigm). Mean heat pain that corresponded to 150 points on the 0- to 200-point eVAS was 46.4°C (SD 0.8°C) and 47.4°C (SD 0.8°C) for 175/200 points respectively. Mean sound stimulation parameters for 150/200 and 175/200 eVAS ratings were 90.8dB (SD 8.0dB) and 95.0dB (SD 7.5dB), respectively.

#### 2.1.4. Experimental paradigm

A TCE paradigm with three successive periods (T1-T2-T3) consisting of an offset trial (OT) and a constant trial (CT) was used [67]. CTs were administered for a duration of 35 seconds, during which continuous heat or sound stimulation was applied at a calibrated intensity corresponding to 150/200 on the eVAS. Since there is no recommended standard to calculate TCE effects in the current literature [67], this method was chosen to not be at risk of overestimating TCE effects. TCE effects were quantified as the difference between OTs and CTs during T3, allowing to control for potential adaptation or sensitization effects unrelated to TCE. The OTs consisted of an intensity corresponding to 150/200 (T1) applied for ten seconds, followed by an intensity of 175/200 (T2) for ten seconds and then decreased back to the stimulation intensity of T1 for 15 seconds (T3). A duration of 15 seconds for T3 was chosen to incorporate enough time to fully capture the temporal dynamics of the offset effect [22,82] while also not unnecessarily prolonging the trial duration. Rise and fall rates were kept constant (100°C/s). Each trial was performed four times (4× OT, 4× CT) with a break of 2 min in between the trials. The order of trials was pseudorandomized in a counterbalanced manner. These eight trials were completed once for each modality, with modality order randomized across participants; e.g. half of the participants received auditory stimulation first followed by thermal stimulation, whereas the other half completed the modalities in the reverse order. Participants continuously rated the experienced pain intensity throughout each trial and were instructed to attend carefully and indicate even very subtle sensations or changes. This procedure was repeated once for each modality, resulting in a total of 16 trials per participant. All trials were completed for one modality before the other modality was tested. This transition was preceded by a 5-min interval of no stimulation to reduce possible carry-over effects. For an overview of the procedure, consult Figure 1.

**Figure 1.**
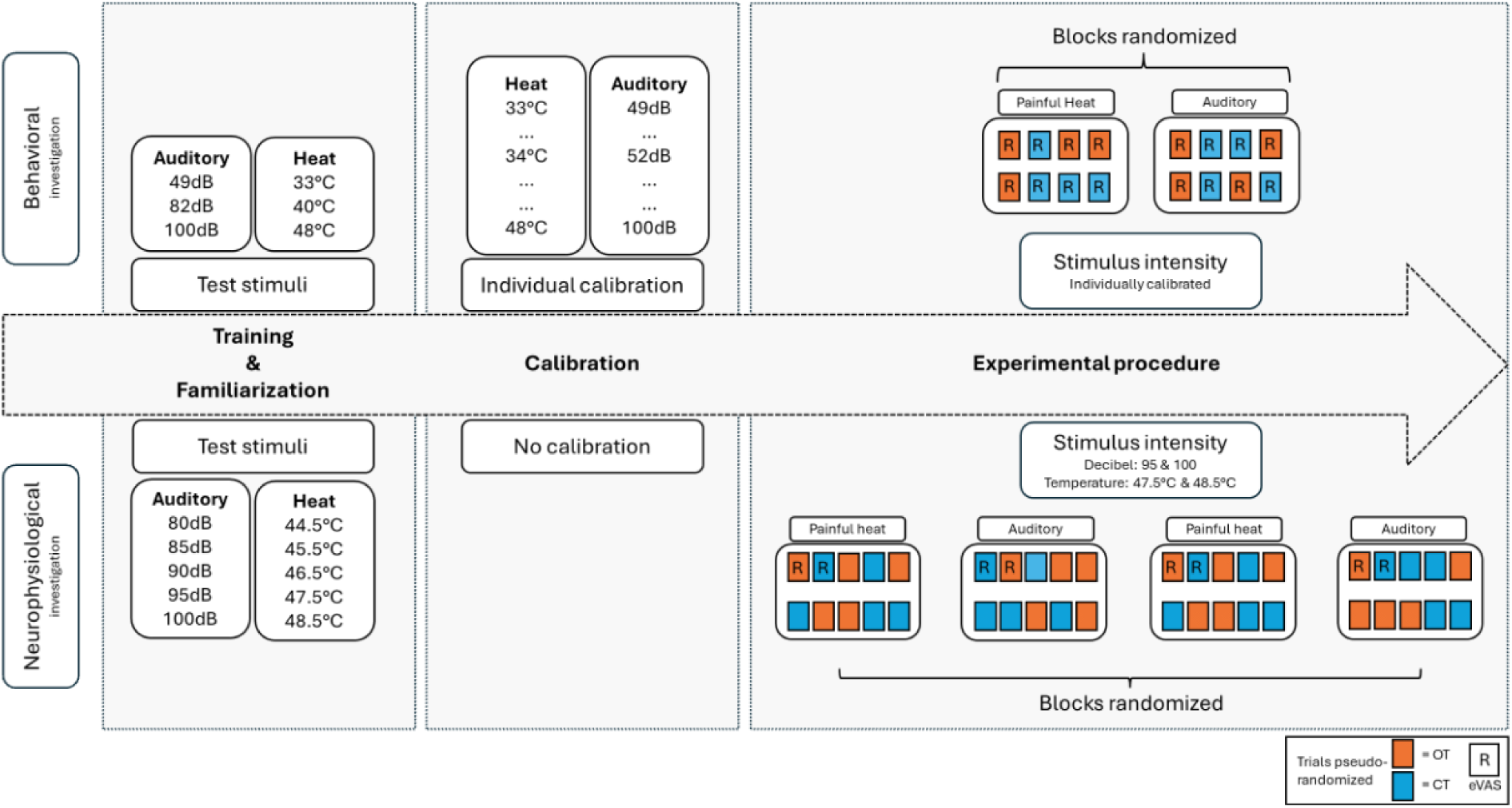
Study design. Schematic representation of the study design of both experiments. Experiment 1 (top of the figure): After preparation, participants underwent a training & familiarization phase to accustom to stimuli and the operation of the electronic visual analogue scale (eVAS). The stimulus intensities were individually calibrated. The experimental procedure was split into two blocks (1 per modality) consisting of 8 trials with offset (OT) or constant (CT) trials in pseudorandomized sequence. In each trial, participants rated their sensory experience using the eVAS. A two-minute pause was conducted after each trial and a five-minute break was conducted after switching to the other modality. In Experiment 2 (bottom of the figure) fixed stimulus intensities were chosen instead of individually calibrating. A block of five test stimuli (per modality) was added instead, so participants could practice eVAS ratings and familiarize to the stimulus intensities. The main experimental procedure was split into four blocks (2 per modality) consisting of 10 trials with offset (OT) or constant (CT) trials in a pseudorandomized sequence. Each trial was followed by a two-minute break, and after each modality block participants had a five-minute break. The first two trials of each block were fixed to include an OT and CT (being randomized in order) and were used to collect the behavioral response. The remaining 8 trials did not include any eVAS ratings, only collecting neurophysiological responses.

### 2.2. Experiment 2: Neurophysiological expression underlying TCE in auditory and thermal stimulation

The aim of this experiment was to exploratively investigate neural and autonomic nervous system responses using a similar stimulation paradigm and the same modalities as explained above. For this, EEG data, pupillometry data and behavioral ratings were collected. The study was approved by the Ethics Committee of the University of Luebeck (2024-438) and conducted in accordance with the Declaration of Helsinki. Again, the methodology was preregistered on the Open Science Framework (OSF; https://osf.io/v37mp).

#### 2.2.1. Participants

A total of 29 healthy participants (sex = 19 females; age = 24.6, SD = 5.7) were used for analysis. In- and Exclusion criteria were similar for both experiments. None of the volunteers that participated in the behaviorally focused experiment were recruited for the second experiment.

#### 2.2.2. Equipment

The same stimulation equipment as in the previous experiment was used for all procedures. Behavioral data were collected using an eVAS similar in style and anchors, but the visual presentation, data collection and stimulus control was achieved using MATLAB (The Mathworks Inc., 2024). For visual presentation purposes, the psychophysics toolbox [4] was used. The EEG was recorded at 24 passive scalp electrodes (SMARTING, mBrainTrain, Belgrade, Serbia) at a sampling rate of 500 Hz (DC to 250 Hz bandwidth), referenced against electrode FCz. Electrode impedances were kept below 10 kΩ. The amplifier was attached to the EEG cap (Easycap, Herrsching, Germany) and the EEG data were transmitted via Bluetooth to a nearby computer, which recorded the data using the Smarting Streamer (Version 3.4.2). For pupillometry, a Tobii X3-120 Eye Tracker (Tobii Technology Inc., Stockholm, Sweden) was used with sampling rate of 120Hz. Pupillometry data were directly recorded via MATLAB.

#### 2.2.3. Familiarization procedure

Contrary to the first experiment, stimulus intensity was not individually calibrated. This approach was chosen due to the robust effects seen in the first experiment. Furthermore, the application of constant stimulation intensities ensures uniform stimulus input across all participants, a feature that assumes greater significance in the context of neurophysiological investigations [1]. The stimulus intensities were derived from mean values of the stimulation parameters of the behavioral experiment. The initially derived stimulation parameters from the first experiment that were used for thermal stimulation led to major pain habituation effects resulting in close to zero pain felt by the participants after a few trials. Due to this, the temperature was increased by 1.5°C (initially 46°C and 47°C) to reduce the observed habituation effects. The participants (n = 11) that received the stimulation protocol prior to this change were excluded from the analysis. This was implemented as a one-time adjustment to the stimulation parameters. Importantly, this decision was not based on habituation to painful stimulation per se; participants showing substantial adaptation after the temperature increase were retained in subsequent data collection and included in both the study sample and statistical analyses. A visual depiction of the eVAS ratings of the excluded participants is provided in the Supporting Information (S10 Fig.) In order to account for the missing practice using the eVAS, the participants underwent a brief familiarization procedure. This procedure comprised five stimuli for each modality in ascending order, with each stimulus being presented for 10 seconds, followed by a 10 second pause (matching the calibration procedure from the first experiment). Thermal stimulation started at 44.5°C and increased 1°C every step with the last stimulus having an intensity of 48.5°C. Auditory stimulation started at an intensity of 80dB, increasing in 5dB steps and finishing with a stimulus of 100dB.

#### 2.2.4. Experimental paradigm

The TCE paradigm matched the behavioral protocol but used fixed stimulation intensities of 47.5°C or 95dB for T1, 48.5°C or 100dB for T2 and 47.5°C or 95dB for T3, respectively. The respective modality was tested in a “stimulus block” consisting of ten trials (OTs and CTs). Each modality was tested twice in total, resulting in a total of 40 trials per participant (2× 10 auditory, 2× 10 thermal stimulation). The first two trials per block were pseudorandomized as either an OT or CT, with the objective of collecting behavioral ratings. Participants were tasked with continuously rating their perceived stimulus intensity. The order of the remaining eight trials was pseudorandomized to an equal number of OTs and CTs. Thus, for an individual trial, participants could not anticipate whether the stimulation intensity of this trial would be constant (CT) or would be raised after 10s (OT). However, participants arguably realized after a few trials of the experiment that if a change in stimulation intensity would occur, this would happen at a fixed point in time (10s). In these trials the participants were instructed to maintain their gaze on a fixation cross, focusing on their sensory experience without moving. Each “modality block” was followed by a 5-min break and then alternated to the other modality, with the starting modality being randomized. EEG preprocessing and analysis.

#### 2.2.5. EEG preprocessing and analysis

The continuous EEG data were high-pass (1 Hz) and low-pass filtered (100 Hz), re-referenced to the average reference across all electrodes, and epoched from –5 to +40s relative to the onset of auditory/thermal stimulation. An independent component analysis (ICA) was used to remove components related to eye-blinks, eye-movements and muscle activity. Remaining artefactual epochs were removed afterwards by visual inspection. All data analyses were carried out in Matlab (R2024b), using custom scripts and the Fieldtrip toolbox [55]. Time-frequency oscillatory power representations of single-trial EEG data were obtained using Fast Fourier Transform (FFT) with multi-tapering (DPSS, discrete prolate spheroidal sequences) for a moving time window (length: 2s; moving in steps of 0.1s through the trial) for frequencies 1–80Hz in steps of 1 Hz with 2Hz spectral smoothing.

#### 2.2.6. Pupillometry preprocessing

Similarly to EEG analysis, Fieldtrip [55] was used to conduct the necessary preprocessing steps for the pupillometry data. First, samples reflecting physiologically implausible pupil changes were identified using a velocity-based criterion: data points whose dilation speed exceeded three standard deviations above the mean velocity within each trial were marked as invalid. Next, short gaps of missing data (< 500 ms) were interpolated using a cubic spline method to reconstruct brief blink-related signal loss, following the recommendations of Sebastiaan Mathôt and Kret and Sjak-Shie [33,42]. Longer gaps were left unaltered to avoid introducing artificial signal. Finally, pupil size data from the left and right eye were averaged to obtain a single mean for subsequent analyses.

### 2.3. Statistical analysis

Statistical analyses were performed using R Studio (RStudio version 2024.04.11 with R version 4.5.0, R Foundation for Statistical Computing, Vienna, Austria) [59] and MATLAB [70]. Parametric data is presented in means (x̄) with standard deviations (SD) and nonparametric data in median (M) with ranges (R) or absolute and relative frequencies. To directly examine potential differences between experiments, we conducted a 2 × 2 × 2 × 3 mixed-design ANOVA with ‘Experiment’ (Experiment 1, Experiment 2) as a between-subject factor and ‘Modality’ (heat, auditory), ‘Trial Type’ (CT, OT), and ‘Time’ (T1, T2, T3) as within-subject factors. In addition, Bayes-Factors were calculated using the bayestestR and BayesFactor packages [39,40,46] to determine whether non-significant findings merely reflected insufficient evidence or provided evidence supporting the absence of an effect. Furthermore, separate 2 × 3 repeated-measures ANOVAs (factors ‘Trial Type’ and ‘Time’) were conducted for each experiment and modality to follow up potential interactions between factors. If there were significant effects, FDR corrected [2,3] post-hoc t-tests were performed. To ensure the robustness of our findings we additionally calculated a non-parametric alternative using the ARTool package [31] which can be accessed in the Supporting Information (S6 Table). The level of significance was set at p < 0.05.

The average eVAS ratings were obtained for each time interval (T1, T2 and T3). The first 5s of T1, T2 and T3 and the last 5s of T3 were not considered in our analysis. This approach was oriented on prior studies to consider the delay in pain response and extract stable pain ratings [38,69]. To analyze the EEG and pupillometry data, the extracted time intervals were shifted to right after onset of the stimulus, since we did not expect any delay in response compared to the behavioral data, resulting in T1 being 0-5s, T2 being 10-15s and T3 being 20-25s. Nevertheless, to ensure the robustness of our findings and to exclude the possibility that relevant effects were masked by differences in temporal segmentation, we conducted supplementary ANOVAs based on time intervals identical to those used in the behavioral analyses. These analyses yielded a comparable pattern of results and are reported in the Supporting Information (S8a and S8b table). In addition to these planned statistical analyses, an exploratory cluster based permutation test [19] was performed, comparing OT and CT during the entire trial time course. This non-parametric approach identifies clusters of contiguous time points exhibiting consistent condition differences and controls for multiple comparisons at the cluster level [41]. The test was conducted using a threshold of p < 0.05 with 10.000 permutations and was conducted as a paired (within-subject) comparison.

To determine whether observed effects in pain are modality specific, an estimation of sample size was done for both experiments. With a combined sample size of n = 62 (across experiments 1 and 2), the statistical power to detect a small-to-medium effect (Cohen’s f = 0.15) for 12 repeated measures – corresponding to the 2 (Modality) × 2 (Trial Type) × 3 (Time) interaction testing the specificity of TCE to nociception – is larger than 95% (G*Power, University of Düsseldorf) [18]. The analysis of (neuro)physiological responses in experiment 2 (n = 29) was exploratory. The goal was to derive a mechanistic understanding that might underly the observed behavioral effects. With a sample size of n = 29 in experiment 2, the statistical power to detect a small-to-medium effect for the difference between time intervals or trial types (Cohen’s d = 0.5 for dependent-samples t-tests) is ∼75% (G*Power, University of Düsseldorf). Lastly, an exploratory Pearson’s correlation analysis was conducted to examine potential associations of TCE effects across modalities. With a combined sample across the two experiments (n = 62), the statistical power to detect a small-to-medium correlation (rho = 0.35) is larger than 80% (G*Power, University of Düsseldorf).

## 3. Results

A total of 62 healthy volunteers were recruited in this study (behavioral investigation: n = 33, behavioral + neurophysiological investigation: n = 29). All participants tolerated the selected stimulus intensities without difficulty and exhibited no signs of adverse events (see S2 and S3 Tables for details). An overview of stimulation parameters, as well as additional analyses for each individual experiment can be accessed in the Materials and Methods Section and the Supporting Information (S5 – S7 Tables).

Before investigating effects of the two experiments separately, a combined ANOVA on participant’s ratings with the factors ‘Experiment’ (1, 2), ‘Modality’ (heat, auditory), ‘Trial Type’ (constant, offset) and ‘Time’ (T1, T2, T3) was performed. The complete ANOVA table is shown in the Supporting Information (S4 Table). A significant main effect of Experiment (F(1, 60) = 36.26, p < 0.001, h_p_^2^ = 0.38) revealed that mean ratings were higher in experiment 1 (mean: 121.5, SEM: 3.61) compared with experiment 2 (mean: 89.8, SEM: 3.85). Furthermore, a main effect of Modality (F(1, 60) = 4.80, p < 0.05, h_p_^2^ = 0.07) indicates differences depending on modality and a significant main effect Time (F (1.91, 114.81) = 161.04, p < 0.001, h_p_^2^ = 0.73) indicates differences across T1-T3. A significant Trial Type × Time interaction (F(1.60, 96.28) = 231.96, p < 0.001, h_p_^2^ = 0.79) indicated a robust TCE effect. In addition, significant Modality × Time (F(1.48, 88.68) = 95.48, p < 0.001, h_p_^2^ = 0.61) and Modality × Trial Type × Time interactions (F(1.56, 93.50) = 52.14, p < 0.001, h_p_^2^ = 0.46) suggested that the magnitude and/or temporal dynamics of TCE differed between modalities. To follow up on these interactions, separate analyses for the two experiments and modalities are reported below.

### 3.1. Experiment 1

#### 3.1.1. Temporal contrast enhancement is robust in heat-induced pain and aversive auditory stimulation

ANOVAs revealed interactions of time x trial type for auditory (F (2, 64) = 43.77, p < 0.001, h_p_^2^ = 0.58) and thermal (F (2, 64) =84.37, p < 0.001, h_p_^2^ = 0.73) stimulation. Comparing trials using false discovery rate (FDR) -corrected post-hoc testing for auditory stimulation revealed no difference between OT and CT during T1 (p = 0.65). However, significantly higher discomfort ratings for OT vs. CT in T2 (p < 0.001) and a reversal of this difference in T3 (p < 0.001) were observed, indicating that a return to the stimulus intensity of T1 induced a significant TCE effect in the interval of interest (T3). Similarly, post hoc testing for thermal stimulation showed no difference between OT and CT during T1(p = 0.49), but significantly higher pain ratings for OT vs. CT in T2 (p < 0.001) and a reversal of this difference in T3 (p < 0.001). These findings demonstrate that TCE – a significant and disproportionate reduction in perceived intensity following a brief decrease in stimulus intensity – occurs for both, painful heat and aversive auditory stimulation.

Additionally, to test temporal filtering within each modality, dependent-samples t-tests were calculated comparing T1 and T3 within each trial type. Auditory stimulation displayed significantly lower discomfort ratings in T1 compared to T3 despite constant stimulation (CT; t (32) = −3.82, p < 0.001, d_z_ = 0.54, M_DIFF_ = −16.14), indicating temporal loudness summation. Whereas no difference between T1 and T3 was found for offset trials (OT; t (32) = 0.27, p = 0.79, d_z_ = 0.04, M_DIFF_ = 1.11). To the contrary, for thermal stimulation, pain ratings decreased from T1 to T3 despite constant stimulation (CT, t (32) = 4.28, p < 0.001, d_z_ = 0.60, M_DIFF_ = 17.81), indicating temporal adaptation. Furthermore, a pronounced decrease in pain ratings from T1 to T3 was observed for offset trials (OT, t (32) = 7.62, p < 0.001, d_z_ = 1.72, M_DIFF_ = 55.50), resulting in a strong TCE effect. Figure 2 provides an Overview of the behavioral results.

Across participants of both experiments, TCE effects (i.e. the OT - CT difference in T3) were not significantly correlated between auditory and thermal stimulation (r = 0.15, p = 0.25; BF_10_ = 0.33), providing evidence in favor of the null hypothesis.

**Figure 2:**
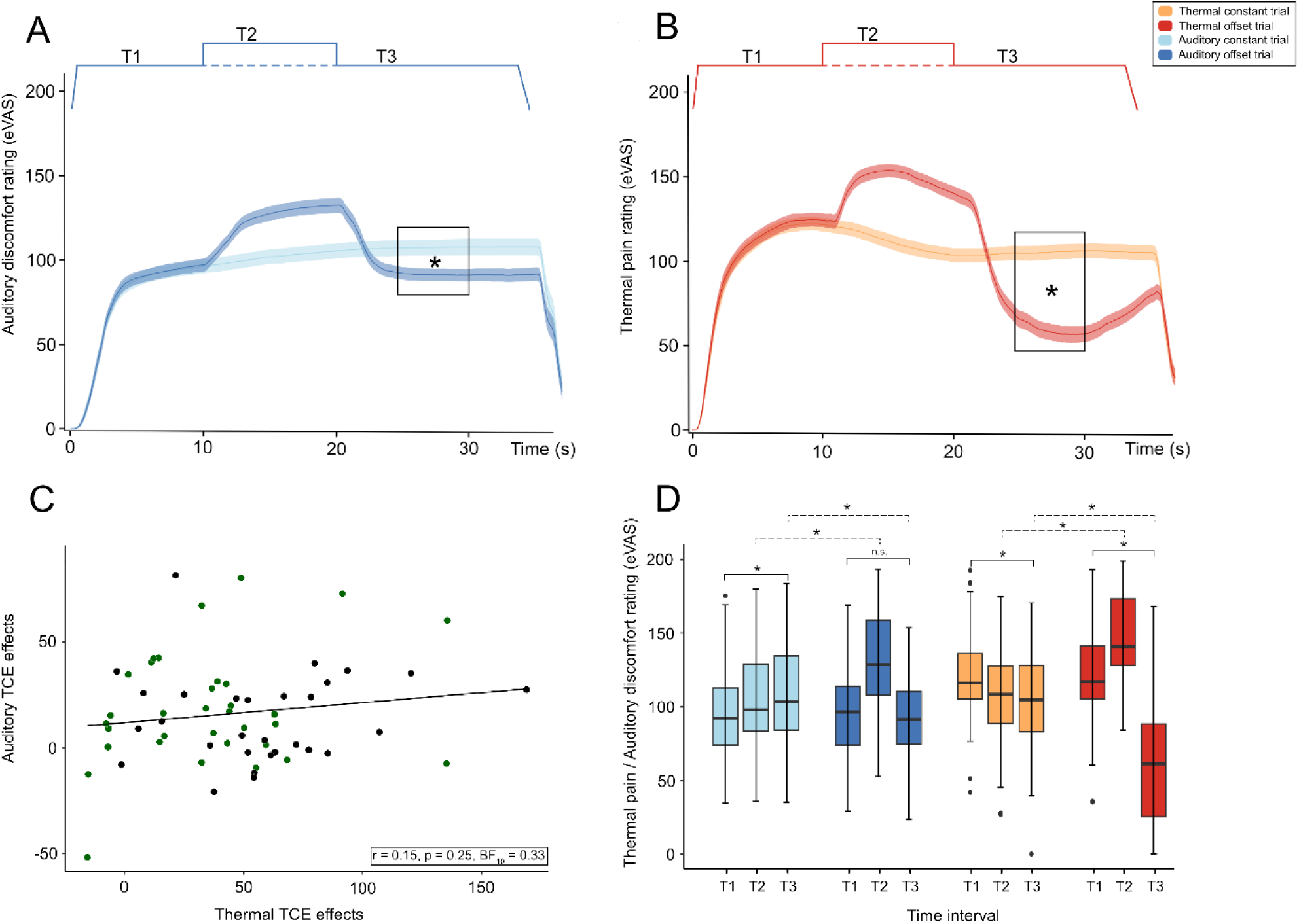
Behavioral Temporal Contrast Enhancement (TCE) response in two modalities. Means and standard errors of the mean (SEM) for behavioral responses are displayed as obtained by an electronic visual analogue scale (eVAS). Auditory stimulation (A) and thermal stimulation (B) both induced significant (*, p < .05) TCE effects in T3 when comparing offset (OT) (darker colors) and constant (CT) (lighter colors) trials. The correlation of TCE effects (C) (CT-OT at T3) (green = behavioral experiment, black = neurophysiological experiment) between both modalities was not significant. Boxplots of eVAS ratings (D) for each of the four conditions split into three relevant time intervals (T1: 5-10s, T2: 15-20s, T3: 25-30s). Significant (p < .05) within-modality comparisons indicated by solid lines, between-modality comparisons indicated by dashed lines.

### 3.2. Experiment 2

The behavioral results were replicated in the second experiment, again indicating a significant interaction between time interval and trial type for auditory (F(2, 56) = 62.10, p < 0.001, h_p_^2^ = 0.69) and thermal (F (2, 56) = 115.22, p < 0.001, h_p_^2^ = 0.80) stimulation, respectively.

#### 3.2.1. Pupil size reflects perception of pain but not auditory discomfort

When investigating the event-related pupil dilation to determine whether TCE is accompanied by modality-dependent autonomic modulation, auditory stimulation displayed a significant main effect of ‘time’ (F (2, 56) = 32.67, p < 0.001, h_p_^2^ = 0.54). FDR corrected post hoc testing revealed significant reduction in the pupil size for T3 vs. T2 (p < 0.001), T3 vs. T1 (p < 0.001) but not for T2 vs. T1 (p = 0.74). This indicates a gradual decrease in pupil size during the trial. For thermal stimulation, there was a significant interaction effect of ‘trial type’ and ‘time’ (F (2, 56) = 64.35, p < 0.001, h_p_^2^ = 0.70). Comparing trial types at different time intervals, resulted in significantly enhanced pupil size for OT vs. CT in T2 (p < 0.001) but not in T1 (p = 0.88) and T3 (p = 0.08). A visual representation of changes in pupil diameter over time for both modalities is displayed in Figure 3.

**Figure 3.**
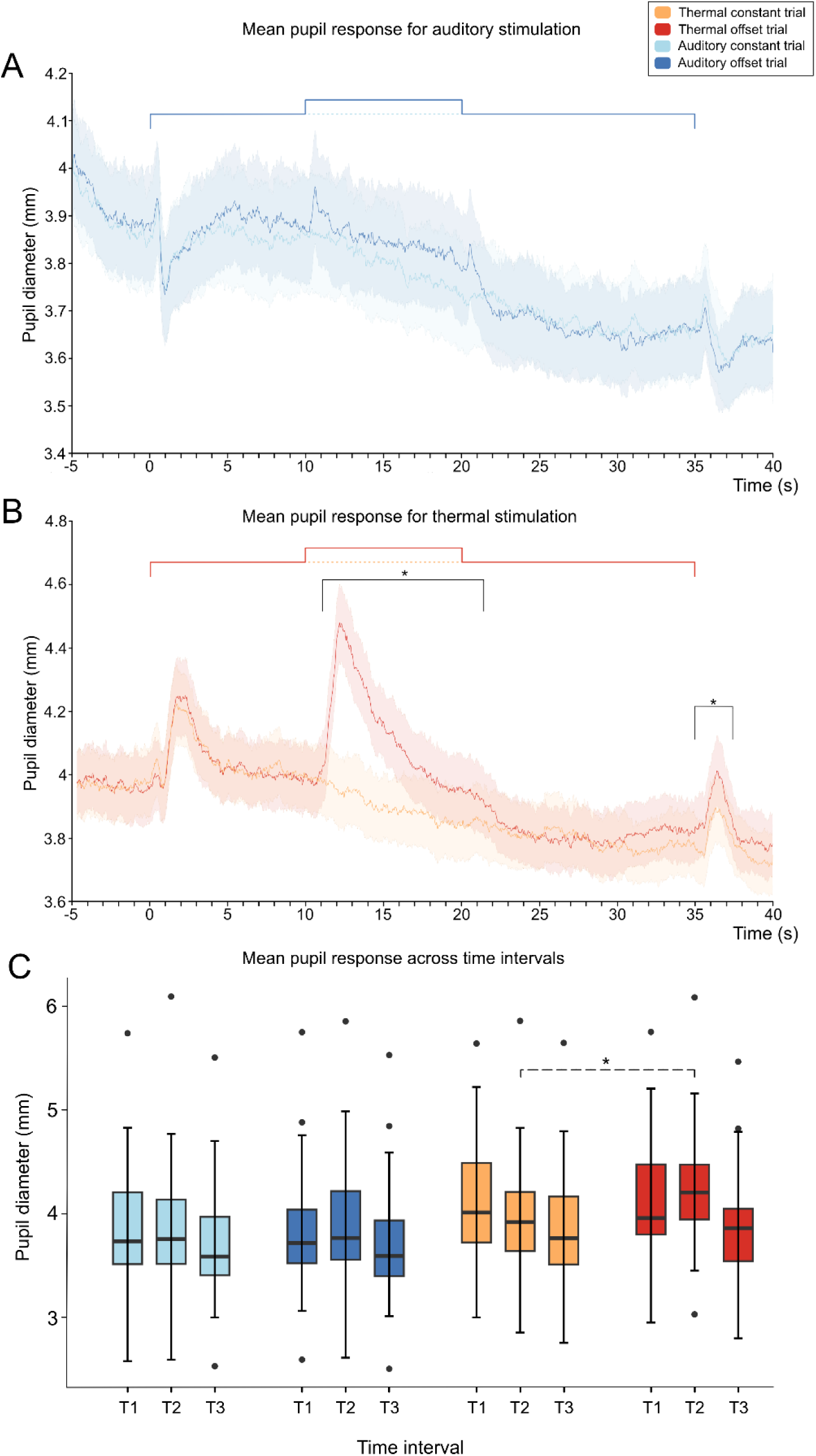
Mean pupil response over time. Mean pupil diameter (mm) over time was measured for auditory (A) and thermal (B) stimulation. Thermal stimulation induced significant changes (*, p < .05; cluster-based permutation test) in the pupil diameter when comparing offset trials (OT, darker colors) with constant trials (CT, lighter colors). No significant clusters could be identified for auditory stimulation, despite visually observable changes to stimulus onset and offset. The timing of the stimulation intensity is displayed as a solid (OT) and dashed (CT) line at the top of the figure and was similar for both modalities. Mean pupil diameter (mm) for each condition (C) split into three time intervals (T1: 0-5s, T2: 15-20s, T3: 20-25s). Significant (p < .05) between-modality comparisons indicated by dashed lines.

#### 3.2.2. Neural alpha power reflects pain sensation and expectation

Conducting a modality-specific 2 (trial type) x 3 (time) ANOVA, revealed a significant main effect of ‘time’ (F (2, 56) = 9.92, p < 0.001, h_p_^2^ = 0.26) for auditory stimulation, while thermal stimulation displayed a significant interaction effect of ‘trial’ and ‘time’ (F (2, 56) = 5.91, p = 0.005, h_p_^2^ = 0.17). Comparing the different time intervals in auditory simulation with FDR corrected post-hoc tests resulted in a significant difference in alpha power between T1 and T2 (p = 0.033), T2 and T3 (p = 0.004), and T3 and T1 (p = 0.004), indicating an overall alpha power increase over time. The interaction effect in thermal stimulation was driven by a significant difference between OT and CT only for T2 (p = 0.002) but not for T1 (p > 0.05) and T3 (p > 0.05). See Figure 4, for a visual representation of alpha power over time.

**Figure 4:**
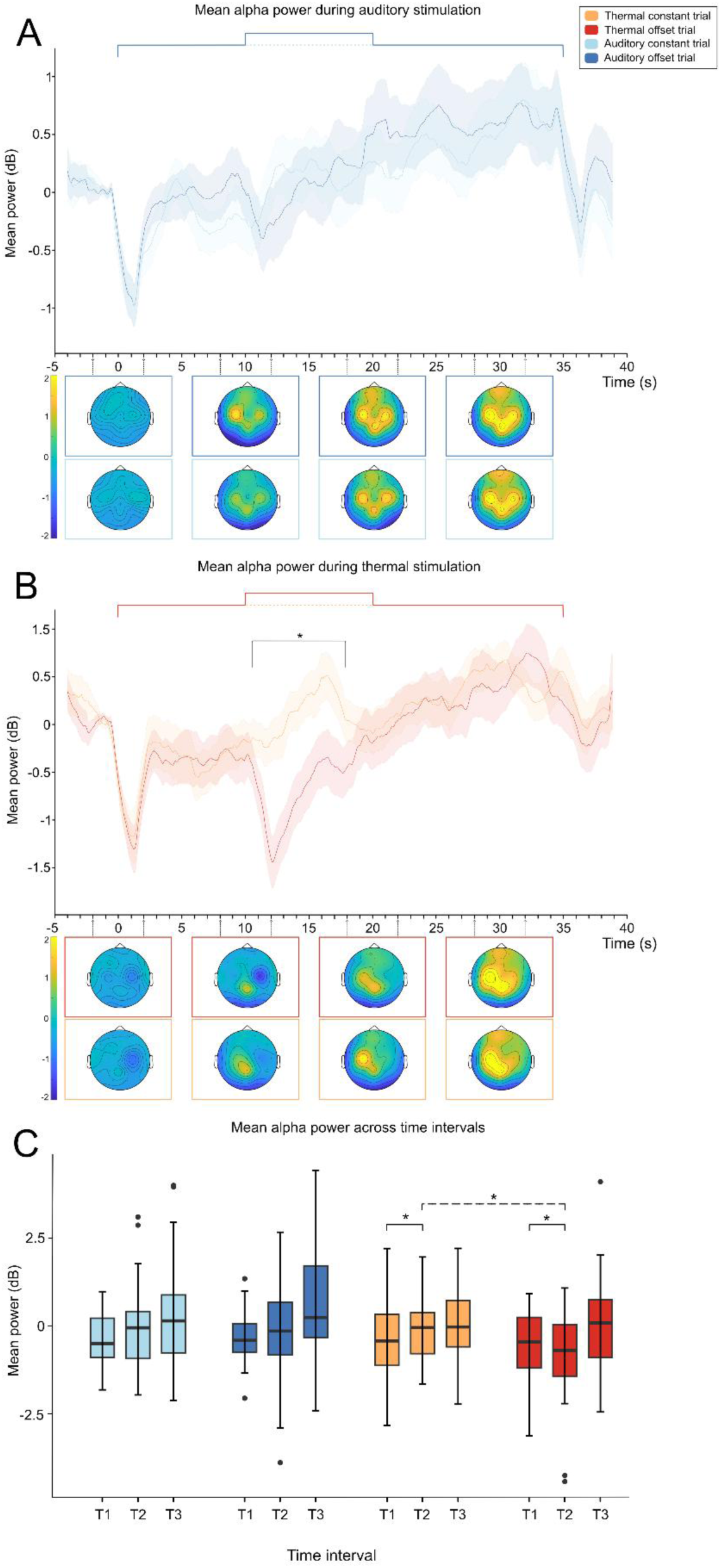
Mean alpha power over time for both modalities. Auditory (**A**) and thermal (**B**) stimulation-induced mean alpha power (7–13 Hz) averaged across centro-occipital electrodes (P7, P3, O1, C3, POz, Pz, CPz, P8, P4, O2, C4). Topographic maps show spatial distributions of alpha power at time points 0, 10, 20 and 30 seconds (color limits: yellow = +2dB, blue = –2dB). Comparing alpha oscillatory power during Offset (OT) (darker colors) and Constant (CT) (lighter colors) trials using cluster-based permutation tests resulted in significant (*, p < .05) differences only during thermal stimulation. Mean (alpha) power for each condition (C) split into the three time intervals of interest (T1: 0-5s, T2: 10-15s, T3: 20-25s). Significant (p < .05) within-modality comparisons indicated by solid lines, between-modality comparisons indicated by dashed lines.

Notably, significant alpha modulation in the T2 interval for thermal stimulation was attributable to two distinct underlying mechanisms reflecting the occurrence and expectation of the stimulus (Figure 5). Post-hoc tests comparing a baseline interval during T1 (5–10 s) to the time interval wherein a cluster-based permutation test indicated a significant difference for OT vs. CT (10.6 - 17.8 s; see Figure 4) revealed the following: First, the increase in thermal stimulation intensity induced an alpha power decrease in OT trials, resulting in significantly lower alpha power compared to T1 (t (28) = −2.61, p = 0.015, d_z_ = 0.48, M_DIFF_ = −0.40). Second, an omission of the expected increase in thermal stimulation intensity induced an alpha power increase in CT trials compared to the baseline interval during T1 (t (28) = 3.34, p = 0.002, d_z_ = 0.62, M_DIFF_ = 0.49). Additionally, a visual representation of the event-related potential can be accessed in the Supporting Information (S11 Fig. and S12 Fig.).

**Figure 5:**
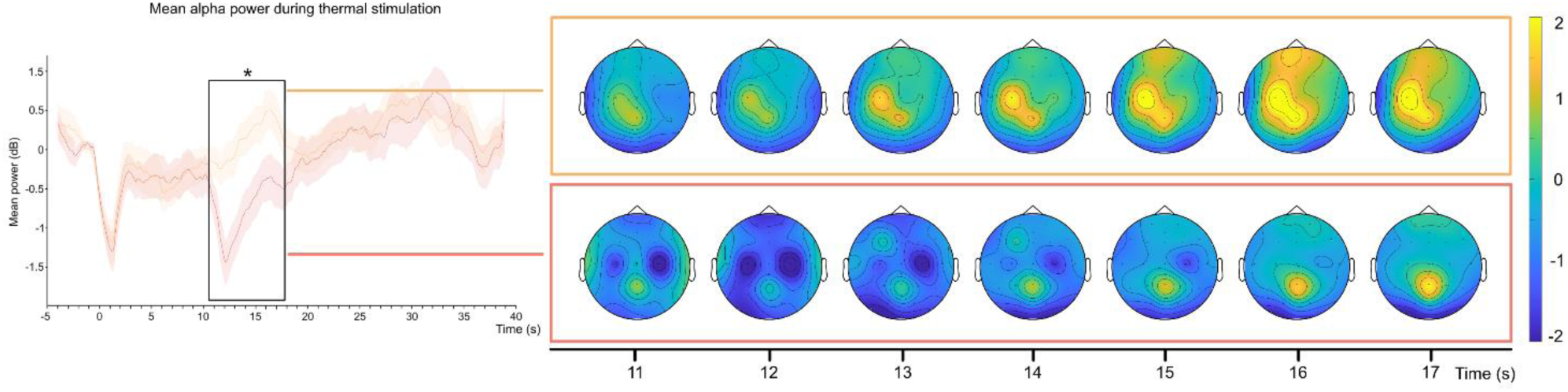
Topographic distribution of alpha power. Stimulation-induced mean alpha power (7–13 Hz) averaged across centro-occipital electrodes (P7, P3, O1, C3, POz, Pz, CPz, P8, P4, O2, C4). Topographic maps show spatial distributions of alpha power at the significant time point (11 sec – 17 sec, one topography per second) of a cluster-based permutation test (* p < .05) comparing offset (OT) and constant (CT) trials (color limits: yellow = +2dB, blue = –2dB).

## 4. Discussion

Temporal filtering operates across sensory modalities, including vision, audition, and pain perception [16,23]. Here, we aimed to delineate behavioral and (neuro-)physiological substrates of temporal filtering and to assess its specificity to pain and unpleasantness. To this end, we employed the phenomenon of TCE, a temporal filtering mechanism thought to underlie offset analgesia [82]. TCE was elicited by pain-inducing heat and aversive sound, possibly indicating a supramodal temporal filtering process with distinct temporal dynamics across modalities. Pupil dilation, indexing autonomic nervous system activation, and alpha-band oscillations, reflecting cortical inhibition, were both modulated by temporal increases in noxious heat but not by auditory stimulation. Surprisingly, these observations did not match the disproportionally large decrease in pain perception that we observed in behavioral TCE ratings, suggesting partially independent mechanisms underlying subjective pain reports and their neurophysiological correlates.

### 4.1. Modality-specific processes underlie temporal contrast enhancement

Consistent with prior evidence [26,67], we were able to successfully induce TCE using thermal stimulation. Noxious heat is known to produce robust TCE effects, indicated by statistically significant decreases in pain sensation following a brief heat offset [26] that far exceed natural adaptation to heat induced pain. To our knowledge, this is the first experiment investigating TCE using non-painful yet aversive auditory stimuli, demonstrating that TCE could reflect a modality-general (i.e., supramodal) mechanism of afferent contrast filtering rather than a nociception-specific process. Conversely, in addition to domain-general effects, we observed modality-specific patterns in subjective ratings, suggesting formal similarities portrayed by contrast effects, with nuanced processing differences between modalities during the stimulation paradigm. When comparing different time intervals (T1 vs. T3) during constant stimulation, we found pain ratings decreasing over time while the opposite was observed for subjective auditory discomfort ratings. We argue that these novel observations represent two different temporal filtering mechanisms. First, adaptation to pain, which is well-known to be induced by tonic heat [21,24]. Second, temporal sound summation, reflecting neural integration of auditory energy over time [6,56].

Comparing ratings between time windows of interest for OTs, we found a typical decrease in perceived pain following heat offset (T3) compared to baseline (T1). In the auditory modality, however, the same contrast was not significant, suggesting a differently constituted TCE effect. Here, TCE was driven by a gradual increase in temporal sound summation during constant stimulation, which was absent during OTs, thereby yielding a comparable overall TCE magnitude.

One possible explanation for the T1 vs. T3 difference, could be adaptation of peripheral nerve fibers induced by heat stimulation, while no equivalent mechanism exists for auditory stimulation [75]. Nevertheless, we would argue that these results could indicate temporal pain inhibition but an absence of such an effect for auditory stimulation. Importantly, this interpretation is further supported by the absence of a correlation between pain- and auditory-evoked TCE effects, which may suggest that the observed effects do not predominantly reflect a single shared CNS mechanism.

### 4.2. Neurophysiological signatures of filtering perception of pain and auditory discomfort

Alongside behavioral responses, we collected objective neuro(physiological) measures (pupillometry and EEG) in a second experiment aiming to better understand the differences between the observed TCE effects for auditory vs. thermal stimulation. Thermal stimulation induced significant differences in pupil size between OT and CT in T2, the interval with the highest stimulus intensity. Reactive increases in pupil diameter to pain-inducing stimuli may reflect autonomic nervous system (ANS) activation, specifically sympathetic arousal associated with heightened awareness and the fight-or-flight response [9,10,13,17,65]. Our observed phasic increases in pupil diameter likely reflect bottom-up processes induced by increases in stimulus temperature and are in line with previous research [10]. However, auditory stimulation did not display such trial type-specific differences, only eliciting significant decreases of pupil diameter over time. This suggests that auditory discomfort did not elicit sizeable modulation of activity in the ANS. Possible explanations for this could be factors such as stimulus intensity or salience, with painful stimulation arguably resulting in a higher perceived threat level compared to auditory stimulation, hence inducing greater changes in pupil diameter.

Consistent with the pupillary dynamics, time–frequency EEG analyses revealed a similar modality-specific pattern: thermal stimulation elicited a significant alpha power reduction during OTs in T2, which was not observed for auditory stimulation. A reduction of alpha power in response to painful stimulation is in line with previous reports of alpha decreases accompanying increased pain perception [25,52,58]. Moreover, when comparing T1 and T2 in OTs, we observed a significant alpha desynchronization, reflecting bottom-up processes associated with increased stimulus intensity. In contrast, CTs exhibited significant alpha synchronization (i.e., an alpha power increase) that may reflect top-down processes, possibly induced by expectation of an increase in painful stimulation. This is in line with prior research observing top-down processes influencing alpha oscillations [57]. For auditory stimulation, we hypothesized similar spatiotemporal patterns of oscillatory activity. However, no significant changes in alpha power were observed for either stimulation pattern. Given the comparable stimulation paradigm employed in both modalities, one could expect that participants would manifest a comparable increase in alpha power, driven by the anticipation of a change in stimulus intensity. Surprisingly, no such effects were observed, which lends further support to the view that the neural modulation and representation of auditory discomfort differ from that of painful stimulation.

Interestingly, we did not observe any neurophysiological changes mirroring the TCE effect that we observed in behavior. Neither pupillometry, nor EEG displayed any significant modulations by trial type (OT vs. CT) during T3. Based on literature showing a robust link between tonic pain and reduced alpha oscillatory power, we hypothesized that the TCE effect would elicit measurable changes in alpha activity — potentially an alpha increase reflecting the experienced relief — as it has been reported for placebo analgesia [28,36]. Contrary, we did not observe any representation of the behavioral TCE effects in neural recordings. One plausible explanation could be that the brain regions responsible for processing or mediating TCE are mainly located subcortically, which would be less accessible to surface EEG. Previous fMRI studies have reported changes to activity in the insula [51,81], structures of the brainstem [15,81], putamen and nucleus accumbens [84] or the spinal cord [63]. Thus, these structures might be more relevant to TCE compared to the more superficially located ones. Additionally, subjective experiences such as changes in pain perception may not always align with objective indices, which could be the case for TCE. Our pupillometry findings provide partial support for this hypothesis, since both modalities displayed temporal filtering as evidenced by decreases in pupil diameter over time. However, these temporal patterns did not correspond with the observed patterns in the subjective domain, where only pain decreased over time, but auditory discomfort increased. To our knowledge, studies investigating TCE employing objective measures besides fMRI are scarce [50,68,73] making it difficult to assess the extent to which subjective and physiological responses to TCE diverge and whether our findings reflect such an incongruence.

We replicated robust behavioral TCE responses across both experiments, using individually calibrated and fixed-stimulus approaches. Cross-modal matching of painful and non-painful stimuli remains inherently challenging, particularly because increasing sound intensity beyond a given threshold was not feasible due to potential risk to participants [8]. Additionally, the lack of individual calibration in the second experiment should be considered when interpreting the results. Nevertheless, fixed-stimulus approaches may offer advantages in neurophysiological settings by reducing procedural burden and participant fatigue, especially when extensive calibration procedures are required [1,49]. Moreover, the absence of substantial evidence for interactions involving experiment type (Experiment 1: calibrated; Experiment 2: non-calibrated) and the other experimental factors suggests that omitting calibration in Experiment 2 did not substantially alter the observed effects. Indeed, we observed modality-specific patterns of temporal dynamics, particularly represented in the behavior of participants. While we cannot fully exclude the possibility that the instruction to report even subtle changes may have lowered the reporting criterion and increased response tendencies, such an effect would be expected to act uniformly across conditions. Importantly, an identical stimulation paradigm was employed for all trials, enabling direct comparison between modalities and providing a common framework for interpreting differences in temporal filtering dynamics. As demonstrated in previous research, TCE has been consistently identified as a robust filtering mechanism in pain. TCE effects can be induced with a variety of stimulus intervals, temperature changes and even repeated instead of tonic stimuli [26,66,67]. Nevertheless, future research could benefit from the use of variable stimulus sequences to further disentangle shared from converging mechanisms that might drive filtering dynamics like TCE across modalities. Additionally, although sample-size estimations were performed, the study may have been underpowered to detect small effects, particularly for exploratory neurophysiological outcomes. Thus, these findings should be viewed as preliminary mechanistic evidence rather than definitive tests of the proposed mechanisms. Future studies should aim to characterize the temporal dynamics of TCE in different sensory modalities without compromising spatial resolution, ideally incorporating approaches that target subcortical regions potentially underlying the effect. Further exploration of pain-specific paradigms, and of potential shared filtering mechanisms across sensory modalities, may provide deeper insight into how such processes shape perceptual experience in our multisensory environment.

## Acknowledgements

The authors have no conflicts of interest. This study is supported by the Deutsche Forschungsgemeinschaft (DFG, German Research Foundation). Data availability statement: Interested individuals may contact the authors directly for access to the data. We thank Anna M. Hagemann for her contribution and assistance during data collection. Additionally, we would like to thank two anonymous reviewers for their valuable feedback and helping us improve the quality of our manuscript.

## References

[1] Adamczyk WM, Szikszay TM, Nahman-Averbuch H, Skalski J, Nastaj J, Gouverneur P, Luedtke K. To Calibrate or not to Calibrate? A Methodological Dilemma in Experimental Pain Research. J Pain 2022;23:1823–1832.

[2] Benjamini Y, Hochberg Y. Controlling the False Discovery Rate: A Practical and Powerful Approach to Multiple Testing. J R Stat Soc Ser B Methodol 1995;57:289–300.

[3] Benjamini Y, Yekutieli D. The Control of the False Discovery Rate in Multiple Testing under Dependency. Ann Stat 2001;29:1165–1188.

[4] Brainard DH, Vision S. The psychophysics toolbox. Spat Vis 1997;10:433–436.

[5] Brooks J, Tracey I. REVIEW: From nociception to pain perception: imaging the spinal and supraspinal pathways. J Anat 2005;207:19–33.

[6] Buus S, Florentine M, Poulsen T. Temporal integration of loudness, loudness discrimination, and the form of the loudness function. J Acoust Soc Am 1997;101:669–680.

[7] Cariani P, Baker JM. Survey of temporal coding of sensory information. Front Comput Neurosci 2025;19:1571109.

[8] Chan HS. Occupational noise exposure; criteria for a recommended standard. National Institute for Occupational Safety and Health. Division of Biomedical and Behavioral Science.;National Institute for Occupational Safety and Health. Education and Information Division.;, 1998 Available: https://stacks.cdc.gov/view/cdc/6376. Accessed 22 July 2023.

[9] Chapman CR, Bradshaw DH, Donaldson GW, Jacobson RC, Nakamura Y. Central noradrenergic mechanisms and the acute stress response during painful stimulation. J Psychopharmacol Oxf Engl 2014;28:1135–1142.

[10] Chapman CR, Oka S, Bradshaw DH, Jacobson RC, Donaldson GW. Phasic pupil dilation response to noxious stimulation in normal volunteers: relationship to brain evoked potentials and pain report. Psychophysiology 1999;36:44–52.

[11] Chen KK, Rolan P, Hutchinson MR, de Zoete RMJ. Reliability of Temporal Summation of Pain in Healthy and Clinical Populations: A Systematic Review and Meta-Analysis. Eur J Pain 2025;29:e70097.

[12] Choi I, Lee J-Y, Lee S-H. Bottom-up and top-down modulation of multisensory integration. Curr Opin Neurobiol 2018;52:115–122.

[13] Chouchou F, Fauchon C, Perchet C, Garcia-Larrea L. An approach to the detection of pain from autonomic and cortical correlates. Clin Neurophysiol 2024;166:152–165.

[14] Clements GM, Gyurkovics M, Low KA, Kramer AF, Beck DM, Fabiani M, Gratton G. Dynamics of alpha suppression index both modality specific and general attention processes. NeuroImage 2023;270:119956.

[15] Derbyshire SWG, Osborn J. Offset analgesia is mediated by activation in the region of the periaqueductal grey and rostral ventromedial medulla. NeuroImage 2009;47:1002–1006.

[16] Di Stefano N, Spence C. Perceiving temporal structure within and between the senses: A multisensory/crossmodal perspective. Atten Percept Psychophys 2025;87:1811–1838.

[17] Eisenach JC, Curry R, Aschenbrenner CA, Coghill RC, Houle TT. Pupil responses and pain ratings to heat stimuli: Reliability and effects of expectations and a conditioning pain stimulus. J Neurosci Methods 2017;279:52–59.

[18] Faul F, Erdfelder E, Lang A-G, Buchner A. G*Power 3: a flexible statistical power analysis program for the social, behavioral, and biomedical sciences. Behav Res Methods 2007;39:175–191.

[19] Gerber EM. permutest. 2025. Available: https://de.mathworks.com/matlabcentral/fileexchange/71737-permutest. Accessed 5 Oct 2025.

[20] Gouverneur P, Li F, Luebke L, Szikszay TM, Roelen SD, Krajewski J, Luedtke K, Grzegorzek M. eVAS: A user-friendly electronic Visual Analogue Scale. J Open Source Softw 2025;10:6876.

[21] Greene LC, Hardy JD. Adaptation of thermal pain in the skin. J Appl Physiol 1962;17:693–696.

[22] Grill JD, Coghill RC. Transient analgesia evoked by noxious stimulus offset. J Neurophysiol 2002;87:2205–2208.

[23] Grothe B, Klump GM. Temporal processing in sensory systems. Curr Opin Neurobiol 2000;10:467–473.

[24] Hashmi JA, Davis KD. Effects of temperature on heat pain adaptation and habituation in men and women. Pain 2010;151:737.

[25] Hauck M, Domnick C, Lorenz J, Gerloff C, Engel AK. Top-down and bottom-up modulation of pain-induced oscillations. Front Hum Neurosci 2015;9:375.

[26] Hermans L, Calders P, Van Oosterwijck J, Verschelde E, Bertel E, Meeus M. An Overview of Offset Analgesia and the Comparison with Conditioned Pain Modulation: A Systematic Literature Review. Pain Physician 2016;19:307–326.

[27] Horing B, Sprenger C, Büchel C. The parietal operculum preferentially encodes heat pain and not salience. PLOS Biol 2019;17:e3000205.

[28] Huneke NTM, Brown CA, Burford E, Watson A, Trujillo-Barreto NJ, El-Deredy W, Jones AKP. Experimental Placebo Analgesia Changes Resting-State Alpha Oscillations. PLoS ONE 2013;8:e78278.

[29] Jensen O, Bonnefond M, VanRullen R. An oscillatory mechanism for prioritizing salient unattended stimuli. Trends Cogn Sci 2012;16:200–206.

[30] Jensen O, Mazaheri A. Shaping Functional Architecture by Oscillatory Alpha Activity: Gating by Inhibition. Front Hum Neurosci 2010;4. doi:10.3389/fnhum.2010.00186.

[31] Kay M, Elkin L, Higgins JJ, Wobbrock JO. mjskay/ARTool: ARTool 0.11.2. 2025. doi:10.5281/zenodo.15192343.

[32] Klimesch W, Sauseng P, Hanslmayr S. EEG alpha oscillations: The inhibition–timing hypothesis. Brain Res Rev 2007;53:63–88.

[33] Kret ME, Sjak-Shie EE. Preprocessing pupil size data: Guidelines and code. Behav Res Methods 2019;51:1336–1342.

[34] Kyle BN, McNeil DW. Autonomic Arousal And Experimentally Induced Pain: A Critical Review of the Literature. Pain Res Manag 2014;19:536859.

[35] Lee I-S, Necka EA, Atlas LY. Distinguishing pain from nociception, salience, and arousal: How autonomic nervous system activity can improve neuroimaging tests of specificity. NeuroImage 2020;204:116254.

[36] Li L, Wang H, Ke X, Liu X, Yuan Y, Zhang D, Xiong D, Qiu Y. Placebo Analgesia Changes Alpha Oscillations Induced by Tonic Muscle Pain: EEG Frequency Analysis Including Data during Pain Evaluation. Front Comput Neurosci 2016;10:45.

[37] Liao H-I, Kidani S, Yoneya M, Kashino M, Furukawa S. Correspondences among pupillary dilation response, subjective salience of sounds, and loudness. Psychon Bull Rev 2016;23:412–425.

[38] Luebke L, von Selle J, Adamczyk WM, Knorr MJ, Carvalho GF, Gouverneur P, Luedtke K, Szikszay TM. Differential Effects of Thermal Stimuli in Eliciting Temporal Contrast Enhancement: A Psychophysical Study. J Pain 2024;25:228–237.

[39] Makowski D, Ben-Shachar MS, Chen SHA, Lüdecke D. Indices of Effect Existence and Significance in the Bayesian Framework. Front Psychol 2019;10:2767.

[40] Makowski D, Ben-Shachar MS, Lüdecke D. bayestestR: Describing Effects and their Uncertainty, Existence and Significance within the Bayesian Framework. J Open Source Softw 2019;4:1541.

[41] Maris E, Oostenveld R. Nonparametric statistical testing of EEG- and MEG-data. J Neurosci Methods 2007;164:177–190.

[42] Mathôt S, Vilotijević A. Methods in cognitive pupillometry: Design, preprocessing, and statistical analysis. Behav Res Methods 2023;55:3055–3077.

[43] Mauk MD, Buonomano DV. The neural basis of temporal processing. Annu Rev Neurosci 2004;27:307–340.

[44] McCracken LM. Pain Vigilance and Awareness Questionnaire (PVAQ). APA PsycTests 1997.

[45] Meyer K, Sprott H, Mannion AF. Cross-cultural adaptation, reliability, and validity of the German version of the Pain Catastrophizing Scale. J Psychosom Res 2008;64:469–478.

[46] Morey RD, Rouder JN, Jamil T, Urbanek S, Forner K, Ly A. BayesFactor: Computation of Bayes Factors for Common Designs. 2026. Available: https://cran.r-project.org/web/packages/BayesFactor/index.html. Accessed 7 Apr 2026.

[47] Morin C, Bushnell MC. Temporal and qualitative properties of cold pain and heat pain: a psychophysical study. Pain 1998;74:67–73.

[48] Mouraux A, Iannetti GD. Nociceptive laser-evoked brain potentials do not reflect nociceptive-specific neural activity. J Neurophysiol 2009;101:3258–3269.

[49] Nahman-Averbuch H, Coghill RC. Pain-autonomic relationships: implications for experimental design and the search for an “objective marker” for pain. Pain 2017;158:2064–2065.

[50] Nahman-Averbuch H, Dayan L, Sprecher E, Hochberg U, Brill S, Yarnitsky D, Jacob G. Sex differences in the relationships between parasympathetic activity and pain modulation. Physiol Behav 2016;154:40–48.

[51] Nahman-Averbuch H, Martucci KT, Granovsky Y, Weissman-Fogel I, Yarnitsky D, Coghill RC. Distinct brain mechanisms support spatial vs temporal filtering of nociceptive information. Pain 2014;155:2491–2501.

[52] Nickel MM, Ta Dinh S, May ES, Tiemann L, Hohn VD, Gross J, Ploner M. Neural oscillations and connectivity characterizing the state of tonic experimental pain in humans. Hum Brain Mapp 2020;41:17–29.

[53] Nir R-R, Sinai A, Moont R, Harari E, Yarnitsky D. Tonic pain and continuous EEG: prediction of subjective pain perception by alpha-1 power during stimulation and at rest. Clin Neurophysiol Off J Int Fed Clin Neurophysiol 2012;123:605–612.

[54] Noppeney U. Perceptual Inference, Learning, and Attention in a Multisensory World. Annu Rev Neurosci 2021;44:449–473.

[55] Oostenveld R, Fries P, Maris E, Schoffelen J-M. FieldTrip: Open Source Software for Advanced Analysis of MEG, EEG, and Invasive Electrophysiological Data. Comput Intell Neurosci 2011;2011:156869.

[56] Pedersen CB, Salomon G. Temporal integration of acoustic energy. Acta Otolaryngol (Stockh) 1977;83:417–423.

[57] Peng W, Babiloni C, Mao Y, Hu Y. Subjective pain perception mediated by alpha rhythms. Biol Psychol 2015;109:141–150.

[58] Peng W, Hu L, Zhang Z, Hu Y. Changes of Spontaneous Oscillatory Activity to Tonic Heat Pain. PLOS ONE 2014;9:e91052.

[59] R Core Team. R: A Language and Environment for Statistical Computing. 2024. Available: https://www.R-project.org/.

[60] Rohenkohl G, Nobre AC. Alpha Oscillations Related to Anticipatory Attention Follow Temporal Expectations. J Neurosci 2011;31:14076–14084.

[61] Sathian K, Lacey S. Cross-Modal Interactions of the Tactile System. Curr Dir Psychol Sci 2022. doi:10.1177/09637214221101877.

[62] Schneider D, Herbst SK, Klatt L-I, Wöstmann M. Target enhancement or distractor suppression? Functionally distinct alpha oscillations form the basis of attention. Eur J Neurosci 2022;55:3256–3265.

[63] Sprenger C, Stenmans P, Tinnermann A, Büchel C. Evidence for a spinal involvement in temporal pain contrast enhancement. NeuroImage 2018;183:788–799.

[64] Strauß A, Wöstmann M, Obleser J. Cortical alpha oscillations as a tool for auditory selective inhibition. Front Hum Neurosci 2014;8. doi:10.3389/fnhum.2014.00350.

[65] Szabadi E. Modulation of physiological reflexes by pain: role of the locus coeruleus. Front Integr Neurosci 2012;6. doi:10.3389/fnint.2012.00094.

[66] Szikszay TM, Adamczyk WM, Lévénez JLM, Gouverneur P, Luedtke K. Temporal properties of pain contrast enhancement using repetitive stimulation. Eur J Pain 2022;26:1437–1447.

[67] Szikszay TM, Adamczyk WM, Luedtke K. The Magnitude of Offset Analgesia as a Measure of Endogenous Pain Modulation in Healthy Participants and Patients With Chronic Pain: A Systematic Review and Meta-Analysis. Clin J Pain 2019;35:189.

[68] Szikszay TM, Adamczyk WM, Panskus J, Heimes L, David C, Gouverneur P, Luedtke K. Psychological mechanisms of offset analgesia: The effect of expectancy manipulation. PloS One 2023;18:e0280579.

[69] Szikszay TM, Melz N, Von Glasenapp B, Adamczyk WM, Luedtke K. Effects of stimulation area and temperature rates on offset analgesia. PAIN Rep 2022;7:e1043.

[70] The Mathworks Inc. MATLAB version: 24.1.0.2578822 (R2024a). 2024. Available: https://www.mathworks.com.

[71] Thermal stimulator. QST-Lab n.d. Available: https://www.qst-lab.eu/tcs-technical-description. Accessed 23 Apr 2026.

[72] Valentini E, Halder S, McInnerney D, Cooke J, Gyimes IL, Romei V. Assessing the specificity of the relationship between brain alpha oscillations and tonic pain. NeuroImage 2022;255:119143.

[73] Van Den Houte M, Van Oudenhove L, Bogaerts K, Van Diest I, Van den Bergh O. Endogenous Pain Modulation: Association with Resting Heart Rate Variability and Negative Affectivity. Pain Med 2018;19:1587–1596.

[74] Wang C-A, Munoz DP. A circuit for pupil orienting responses: implications for cognitive modulation of pupil size. Curr Opin Neurobiol 2015;33:134–140.

[75] Willmore BDB, King AJ. Adaptation in auditory processing. Physiol Rev 2023;103:1025–1058.

[76] Wilsch A, Henry MJ, Herrmann B, Maess B, Obleser J. Alpha Oscillatory Dynamics Index Temporal Expectation Benefits in Working Memory. Cereb Cortex 2015;25:1938–1946.

[77] Wöstmann M, Alavash M, Obleser J. Alpha Oscillations in the Human Brain Implement Distractor Suppression Independent of Target Selection. J Neurosci 2019;39:9797–9805.

[78] Wöstmann M, Herrmann B, Maess B, Obleser J. Spatiotemporal dynamics of auditory attention synchronize with speech. Proc Natl Acad Sci 2016;113:3873–3878.

[79] Wöstmann M, Herrmann B, Wilsch A, Obleser J. Neural Alpha Dynamics in Younger and Older Listeners Reflect Acoustic Challenges and Predictive Benefits. J Neurosci 2015;35:1458–1467.

[80] Wöstmann M, Maess B, Obleser J. Orienting auditory attention in time: Lateralized alpha power reflects spatio-temporal filtering. NeuroImage 2021;228:117711.

[81] Yelle MD, Oshiro Y, Kraft RA, Coghill RC. Temporal filtering of nociceptive information by dynamic activation of endogenous pain modulatory systems. J Neurosci Off J Soc Neurosci 2009;29:10264–10271.

[82] Yelle MD, Rogers JM, Coghill RC. Offset analgesia: A temporal contrast mechanism for nociceptive information. Pain 2008;134:174–186.

[83] Zekveld AA, Koelewijn T, Kramer SE. The Pupil Dilation Response to Auditory Stimuli: Current State of Knowledge. Trends Hear 2018;22:2331216518777174.

[84] Zhang S, Li T, Kobinata H, Ikeda E, Ota T, Kurata J. Attenuation of offset analgesia is associated with suppression of descending pain modulatory and reward systems in patients with chronic pain. Mol Pain 2018;14:1744806918767512.

[85] Zhu Y, Nachtrab G, Keyes PC, Allen WE, Luo L, Chen X. Dynamic salience processing in paraventricular thalamus gates associative learning. Science 2018;362:423–429.

[86] Zimmer K, Ellermeier W. Eine deutsche Version der Lärmempfindlichkeitsskala von Weinstein. Z Für Lärmbekämpfung 1997;44:107–110.

