## Supporting_Information for "Temporal Contrast Enhancement in Auditory and Nociceptive Processing"

for the article

### S1 Fig. Calibration procedure in the behavioral investigation

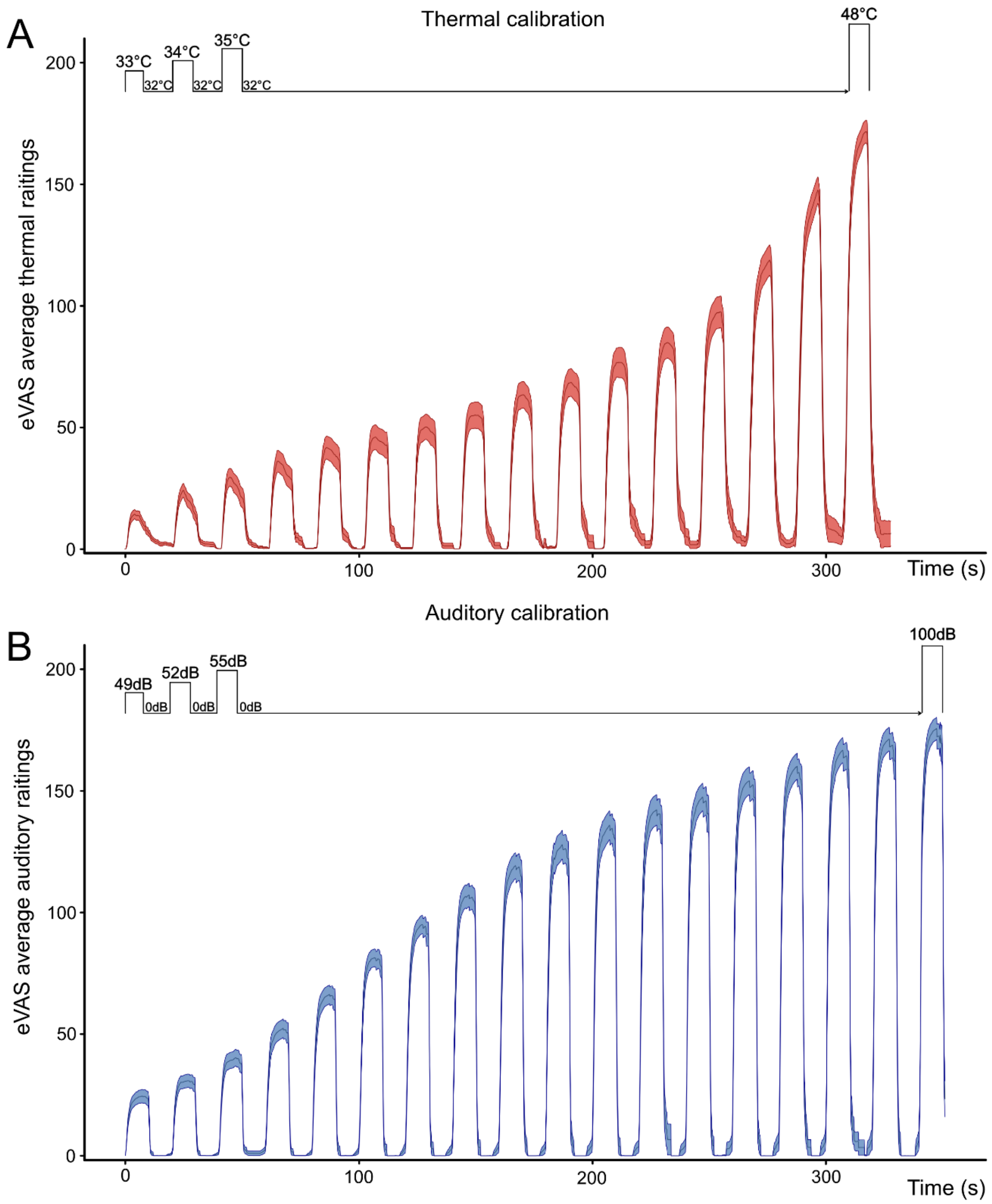


**Calibration Procedure.** Behavioral ratings (behavioral experiment, n = 33) were collected using an electronical visual analogue scale (eVAS). The calibration procedure followed a staircase approach and was repeated twice in thermal (**A**) and auditory (**B**) stimulation. Thermal stimulation started at 33°C, increasing by 1°C per step with each stimulus being applied for 10 seconds. and then returning to baseline temperature (32°C) for 10 seconds. The calibration procedure ended after the final stimulation intensity of 48°C. Auditory stimulation started at 49dB, increasing 3dB per step and followed the same stimulation pattern of 10 seconds stimulus followed by 10 seconds of no stimulus (0dB), ending at a final stimulus intensity of 100dB.

**S2 Table. Participant characteristics in the behavioral investigation**

| **Characteristics, n = 33** | | |
|---|---|---|
| **Age** | Years, x̄ (SD) | 23.7 (6.0) |
| **Height** | Cm, x̄ (SD) | 173.6 (8.7) |
| **Weight** | Kg, x̄ (SD) | 71.9 (18.1) |
| **Female** | N (%) | 22 (66.7) |
| **Right-handed** | N (%) | 33 (100) |
| **Fear of heat pain** | VAS, x̄ (SD) | 38.3 (25.9) |
| **Fear of noise** | VAS, x̄ (SD) | 36.4 (25.9) |
| **Questionnaires** | | |
| **PCS** | M (R) | 15 (0-37) |
| **PVAQ** | | 32 (17-52) |
| **WNSS** | | 52 (14-85) |

*PCS, Pain Sensitivity Questionnaire (range 0-52); PVAQ, Pain and vigilance awareness questionnaire (range 0-80); WSQ, Weinstein Noise Sensitivity Scale (WNSS) (range 0-105); x̄, mean; SD, standard deviation; M, median; R, range.*

**S3 Table. Participant characteristics in the neurophysiological investigation**

| **Characteristics, n = 29** | | |
|---|---|---|
| **Age** | Years, x̄ (SD) | 24.6 (5.7) |
| **Height** | Cm, x̄ (SD) | 172.6 (9.9) |
| **Weight** | Kg, x̄ (SD) | 68.6 (12.5) |
| **Female** | N (%) | 19 (65.5) |
| **Right-handed** | N (%) | 26 (89.7) |
| **Fear of heat pain** | VAS, x̄ (SD) | 41.7 (22.2) |
| **Fear of noise** | VAS, x̄ (SD) | 42.5 (20.9) |
| **Questionnaires** | | |
| **PCS** | M (R) | 14 (0-30) |
| **PVAQ** | | 39 (22-58) |
| **WNSS** | | 60 (27-86) |

*PCS, Pain Sensitivity Questionnaire; PVAQ, Pain and vigilance awareness questionnaire; WNSS, Weinstein Noise Sensitivity Scale; x̄, mean; SD, standard deviation; M, median; R, range.*

**S4 Table. Combined repeated measures ANOVA**

| Row | num Df | den DF | MSE | F | ges | Pr(>F) | log_BF |
|---|---|---|---|---|---|---|---|
| **Experiment** | 1.00 | 60.00 | 5.162.32 | 36.26 | 0.23 | 0.00 | 11.29 |
| **Modality** | 1.00 | 60.00 | 1.947.07 | 4.80 | 0.01 | 0.03 | 4.97 |
| **Experiment:Modality** | 1.00 | 60.00 | 1.947.07 | 0.42 | 0.00 | 0.52 | -1.46 |
| **Trialtype** | 1.00 | 60.00 | 461.06 | 0.78 | 0.00 | 0.38 | -2.20 |
| **Experiment:Trialtype** | 1.00 | 60.00 | 461.06 | 0.08 | 0.00 | 0.78 | -2.09 |
| **Time** | 1.91 | 114.81 | 400.47 | 161.04 | 0.17 | 0.00 | 66.43 |
| **Experiment:Time** | 1.91 | 114.81 | 400.47 | 0.54 | 0.00 | 0.58 | -3.20 |
| **Modality:Trialtype** | 1.00 | 60.00 | 386.23 | 3.08 | 0.00 | 0.08 | -1.14 |
| **Experiment:Modality:Trialtype** | 1.00 | 60.00 | 386.23 | 3.20 | 0.00 | 0.08 | -0.56 |
| **Modality:Time** | 1.48 | 88.68 | 441.11 | 95.48 | 0.09 | 0.00 | 53.14 |
| **Experiment:Modality:Time** | 1.48 | 88.68 | 441.11 | 0.80 | 0.00 | 0.42 | -2.35 |
| **Trialtype:Time** | 1.60 | 96.28 | 355.91 | 231.96 | 0.18 | 0.00 | 107.04 |
| **Experiment:Trialtype:Time** | 1.60 | 96.28 | 355.91 | 2.56 | 0.00 | 0.09 | -1.25 |
| **Modality:Trialtype:Time** | 1.56 | 93.50 | 202.25 | 52.14 | 0.03 | 0.00 | 12.36 |
| **Experiment:Modality:Trialtype:Time** | 1.56 | 93.50 | 202.25 | 4.31 | 0.00 | 0.02 | -1.11 |

A 2x2x2x3 repeated measures ANOVA of the behavioral data combining both investigations (n = 62) including four levels: ‘Experiment’ (1, 2), ‘Modality’ (heat, auditory), ‘Trial Type’ (Offset trial / Constant trial) and ‘Time’ (T1, T2, T3). Extracted time intervals were T1 (5-10s), T2 (15-20s) and T3 (25-30s). The level of significance was set at $p < 0.05$. MSE = mean squared error; ges = generalized eta squared (effect size); Pr(>F) = p-value; log_BF = ln(Inclusion Bayes Factor).

**S5a Table. Behavioral investigation: Repeated measures ANOVA (auditory modality)**

| Row | num Df | den DF | MSE | F | ges | Pr(>F) |
|---|---|---|---|---|---|---|
| **Trialtype** | 1.00 | 32.00 | 476.35 | 0.33 | 0.00 | 0.57 |
| **Time** | 1.77 | 56.74 | 327.76 | 32.29 | 0.11 | 0.00 |
| **Trialtype:Time** | 1.93 | 61.71 | 177.34 | 43.77 | 0.09 | 0.00 |

A 2x3 repeated measures ANOVA of the behavioral data in the auditory modality collected in the behavioral investigation (n = 33) including two levels: 'trial' (Offset trial / Constant trial) and 'time' (T1, T2, T3). Extracted time intervals were T1 (5-10s), T2 (15-20s) and T3 (25-30s). The level of significance was set at $p < 0.05$. MSE = mean squared error; ges = generalized eta squared (effect size); Pr(>F) = p-value

**S5b Table. Behavioral investigation: Repeated measures ANOVA (painful-heat modality)**

| Row | num Df | den DF | MSE | F | ges | Pr(>F) |
|---|---|---|---|---|---|---|
| **Trialtype** | 1.00 | 32.00 | 444.95 | 0.39 | 0.00 | 0.54 |
| **Time** | 1.47 | 47.14 | 696.37 | 77.98 | 0.32 | 0.00 |
| **Trialtype:Time** | 1.39 | 44.43 | 403.49 | 84.37 | 0.22 | 0.00 |

A 2x3 repeated measures ANOVA of the behavioral data in the painful-heat modality collected in the behavioral investigation (n = 33) including two levels: 'trial' (Offset trial / Constant trial) and 'time' (T1, T2, T3). Extracted time intervals were T1 (5-10s), T2 (15-20s) and T3 (25-30s). The level of significance was set at $p < 0.05$. MSE = mean squared error; ges = generalized eta squared (effect size); Pr(>F) = p-value

**S6 Table. Behavioral investigation: Comparison of parametric and non-parametric tests**

| **Effect** | **F** | **Pr(>F)** | **Method** | **Modality** |
|---|---|---|---|---|
| Trialtype | 0.39 | 0.54 | Parametric | Thermal |
| Time | 77.98 | 0.00 | Parametric | Thermal |
| Trialtype:Time | 84.37 | 0.00 | Parametric | Thermal |
| Trialtype | 0.94 | 0.34 | ART | Thermal |
| Time | 81.35 | 0.00 | ART | Thermal |
| Trialtype:Time | 86.36 | 0.00 | ART | Thermal |
| Trialtype | 0.33 | 0.57 | Parametric | Auditory |
| Time | 32.29 | 0.00 | Parametric | Auditory |
| Trialtype:Time | 43.77 | 0.00 | Parametric | Auditory |
| Trialtype | 0.49 | 0.49 | ART | Auditory |
| Time | 35.41 | 0.00 | ART | Auditory |
| Trialtype:Time | 46.27 | 0.00 | ART | Auditory |

A comparison of a 2x3 repeated measures ANOVA of the behavioral data in both modalities with a non-parametric alternative (Aligned Rank Transform; ART, ARTools package in R, Kay et al., 2025) including two levels: ‘trial’ (Offset trial / Constant trial) and ‘time’ (T1, T2, T3) was conducted. Extracted time intervals were T1 (5-10s), T2 (15-20s) and T3 (25-30s). The level of significance was set at $p < 0.05$. Pr(>F) = p-value.

**S7a Table. Neurophysiological investigation: Repeated measures ANOVA of the behavioral data (auditory modality)**

| Row | num Df | den DF | MSE | F | ges | Pr(>F) |
|---|---|---|---|---|---|---|
| **Trialtype** | 1.00 | 28.00 | 220.99 | 7.26 | 0.01 | 0.01 |
| **Time** | 1.78 | 49.98 | 186.47 | 57.99 | 0.12 | 0.00 |
| **Trialtype:Time** | 1.74 | 48.84 | 131.50 | 62.10 | 0.09 | 0.00 |

A 2x3 repeated measures ANOVA of the behavioral data in the auditory modality collected in the neurophysiological investigation (n = 29) including two levels: ‘trial’ (Offset trial / Constant trial) and ‘time’ (T1, T2, T3). Extracted time intervals were T1 (5-10s), T2 (15-20s) and T3 (25-30s). The level of significance was set at $p < 0.05$. MSE = mean squared error; ges = generalized eta squared (effect size); Pr(>F) = p-value

**S7b Table. Neurophysiological investigation: Repeated measures ANOVA of the behavioral data (painful-heat modality)**

| Row | num Df | den DF | MSE | F | ges | Pr(>F) |
|---|---|---|---|---|---|---|
| **Trialtype** | 1.00 | 28.00 | 541.71 | 1.39 | 0.00 | 0.25 |
| **Time** | 1.81 | 50.64 | 480.80 | 79.56 | 0.31 | 0.00 |
| **Trialtype:Time** | 1.46 | 40.93 | 437.04 | 115.22 | 0.32 | 0.00 |

A 2x3 repeated measures ANOVA of the behavioral data in the painful heat modality collected in the neurophysiological investigation (n = 29) including two levels: ‘trial’ (Offset trial / Constant trial) and ‘time’ (T1, T2, T3). Extracted time intervals were T1 (5-10s), T2 (15-20s) and T3 (25-30s). The level of significance was set at $p < 0.05$. MSE = mean squared error; ges = generalized eta squared (effect size); Pr(>F) = p-value

**S8a Table. Comparison of extracted time intervals: Repeated measures ANOVA of the pupillometry data**

| **Pupillometry** | | | | | | | | |
|---|---|---|---|---|---|---|---|---|
| **Auditory** | | | | | | | | |
| | | **num Df** | **den Df** | **MSE** | **F** | **ges** | **Pr(>F)** | **log_BF** |
| Time interval matching behavioral analysis | Trialtype | 1.00 | 28.00 | 0.01 | 4.33 | 0.00 | 0.05 | -0.22 |
| | Time | 1.44 | 40.37 | 0.02 | 42.84 | 0.02 | 0.00 | 38.55 |
| | Trialtype:Time | 1.58 | 44.27 | 0.01 | 2.87 | 0.00 | 0.08 | -0.87 |
| Time interval shifted (-5s) | Trialtype | 1.00 | 28.00 | 0.01 | 1.57 | 0.00 | 0.22 | -1.02 |
| | Time | 1.92 | 53.73 | 0.01 | 32.67 | 0.01 | 0.00 | 31.26 |
| | Trialtype:Time | 1.84 | 51.43 | 0.00 | 1.23 | 0.00 | 0.30 | -1.83 |
| **Heat** | | | | | | | | |
| | | **num Df** | **den Df** | **MSE** | **F** | **ges** | **Pr(>F)** | **log_BF** |
| Time interval matching behavioral analysis | Trialtype | 1.00 | 28.00 | 0.01 | 10.85 | 0.00 | 0.00 | 1.94 |
| | Time | 1.79 | 50.21 | 0.02 | 47.06 | 0.03 | 0.00 | 38.43 |
| | Trialtype:Time | 1.66 | 46.36 | 0.01 | 26.87 | 0.00 | 0.00 | 8.25 |
| Time interval shifted (-5s) | Trialtype | 1.00 | 28.00 | 0.01 | 51.00 | 0.01 | 0.00 | 9.64 |
| | Time | 1.44 | 40.28 | 0.03 | 43.66 | 0.04 | 0.00 | 38.41 |
| | Trialtype:Time | 1.91 | 53.45 | 0.00 | 64.35 | 0.01 | 0.00 | 14.40 |

A 2x3 repeated measures ANOVA of the pupillometry data in the auditory and painful heat modality including two factors: 'Trial Type' (Offset trial / Constant trial) and 'Time' (T1, T2, T3) comparing two adjacent time intervals. Extracted time intervals were either matching behavioral analysis: T1 (5-10s), T2 (15-20s) and T3 (25-30s); or shifted -5 seconds: T1 (0-5s), T2 (10-15s) and T3 (20-25s). The level of significance was set at $p < 0.05$. MSE = mean squared error; ges = generalized eta squared (effect size); Pr(>F) = p-value; log_BF = ln(Inclusion Bayes Factor).

**S8b Table. Comparison of extracted time intervals: Repeated measures ANOVA of the EEG data**

| **EEG Alpha Power** | | | | | | | | |
|---|---|---|---|---|---|---|---|---|
| **Auditory** | | | | | | | | |
| | | **num Df** | **den Df** | **MSE** | **F** | **ges** | **Pr(>F)** | **log_BF** |
| Time interval matching behavioral analysis | Trialtype | 1.00 | 28.00 | 1.24 | 1.09 | 0.00 | 0.31 | -0.87 |
| | Time | 1.29 | 36.18 | 1.36 | 5.62 | 0.03 | 0.02 | 3.50 |
| | Trialtype:Time | 1.60 | 44.74 | 0.20 | 0.87 | 0.00 | 0.40 | -2.09 |
| Time interval shifted (-5s) | Trialtype | 1.00 | 28.00 | 0.79 | 0.13 | 0.00 | 0.72 | -1.74 |
| | Time | 1.26 | 35.31 | 1.53 | 9.92 | 0.07 | 0.00 | 9.37 |
| | Trialtype:Time | 1.96 | 54.93 | 0.26 | 2.76 | 0.01 | 0.07 | -1.43 |
| **Heat** | | | | | | | | |
| | | **num Df** | **den Df** | **MSE** | **F** | **ges** | **Pr(>F)** | **log_BF** |
| Time interval matching behavioral analysis | Trialtype | 1.00 | 28.00 | 1.76 | 1.34 | 0.01 | 0.26 | -0.22 |
| | Time | 1.51 | 42.40 | 0.62 | 16.09 | 0.05 | 0.00 | 6.24 |
| | Trialtype:Time | 1.95 | 54.73 | 0.36 | 3.54 | 0.01 | 0.04 | -0.87 |
| Time interval shifted (-5s) | Trialtype | 1.00 | 28.00 | 1.38 | 3.58 | 0.02 | 0.07 | 0.93 |
| | Time | 1.48 | 41.46 | 1.12 | 9.24 | 0.06 | 0.00 | 4.90 |
| | Trialtype:Time | 1.75 | 48.90 | 0.54 | 5.91 | 0.02 | 0.01 | 0.50 |

A 2x3 repeated measures ANOVA of the EEG data in the auditory and painful heat modality including two factors: ‘Trial Type‘ (Offset trial / Constant trial) and ‘Time‘ (T1, T2, T3) comparing two adjacent time intervals. Extracted time intervals were either matching behavioral analysis: T1 (5-10s), T2 (15-20s) and T3 (25-30s); or shifted -5 seconds: T1 (0-5s), T2 (10-15s) and T3 (20-25s). The level of significance was set at $p < 0.05$. MSE = mean squared error; ges = generalized eta squared (effect size); Pr(>F) = p-value; log_BF = ln(Inclusion Bayes Factor).

### S9 Fig. Facilitatory vs. inhibitory TCE effects

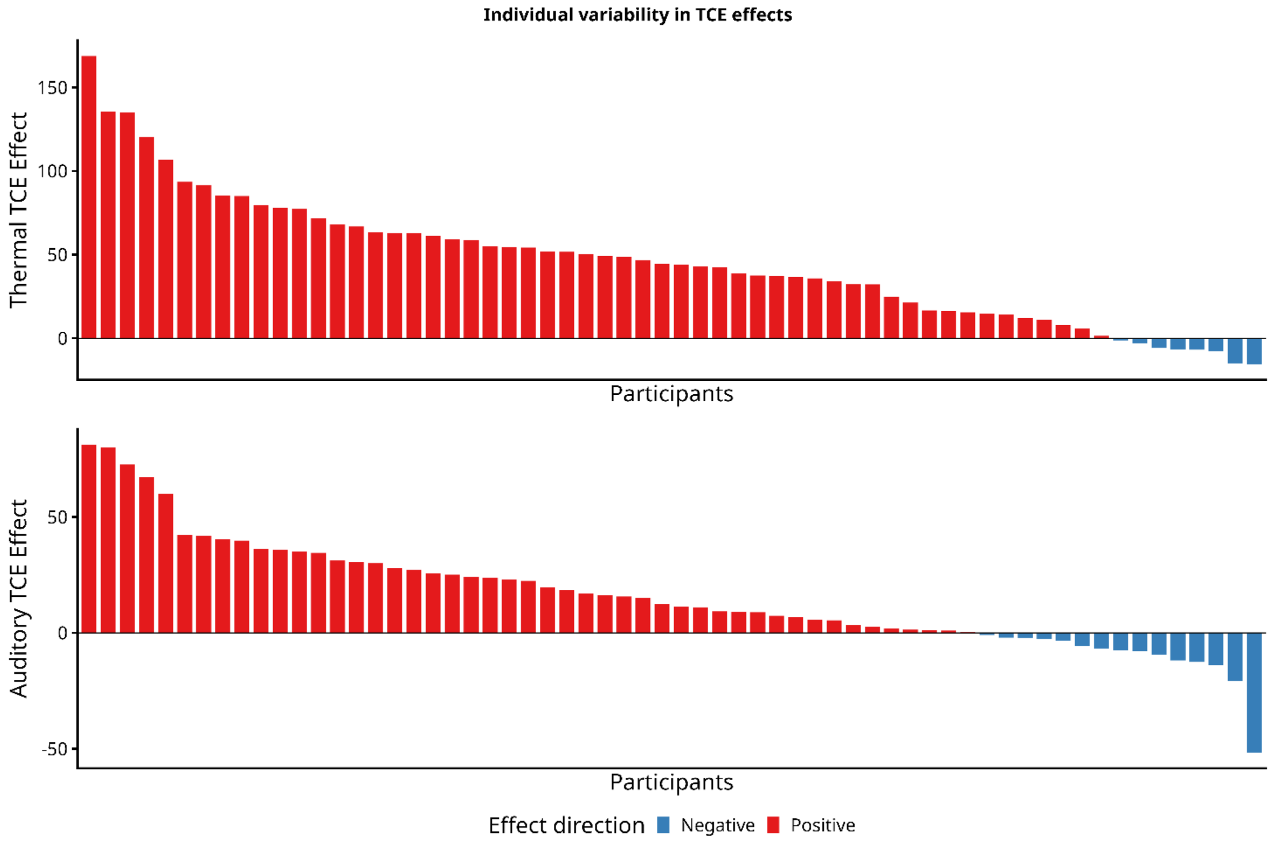


**Facilitatory vs. inhibitory TCE effects**: Individual variability in TCE responses resulting in facilitatory or inhibitory TCE effects. Sorted in order of magnitude across participants for thermal (top) and auditory (bottom) stimulation. TCE effects were calculated by subtracting mean constant trial (CT) values from offset trial (OT) values in the T3 time interval (25-30s). Positive values (red) indicate a present (inhibitory) TCE effect (higher ratings in CT compared to OT trials), whereas negative values (blue) indicate a reversal of the TCE effect resulting in facilitated pain ratings (higher ratings in OT compared to CT trials).

**S10 Fig. Heat pain ratings of excluded participants (n=11)**

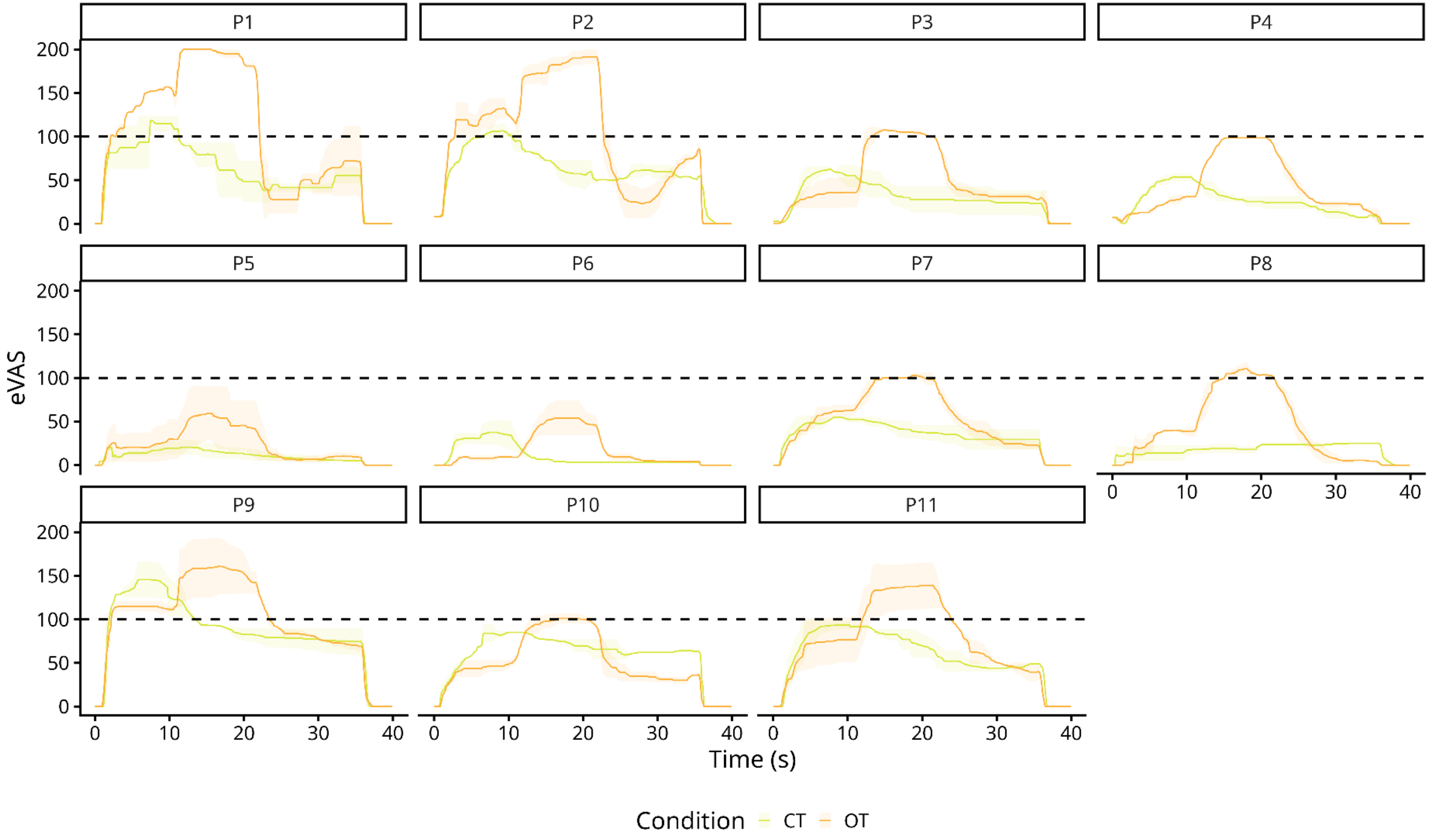


**Heat pain ratings of the excluded participants.** Average ratings (eVAS) during offset trials (OT, orange) and constant trials (CT, green) for heat stimulation. A total of eleven participants were excluded from the analysis due to changes to the experimental paradigm (increase in fixed stimulus intensities of the painful heat modality).

**S11 Fig. Low-pass filtered event-related potential (ERP) for auditory and thermal stimulation**

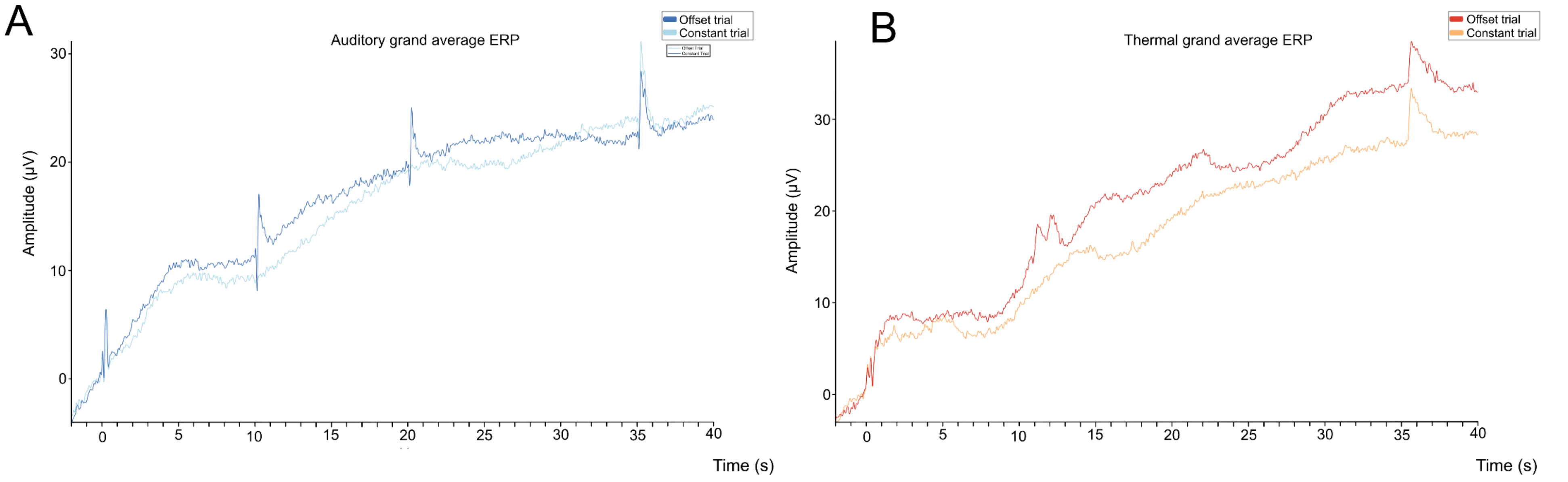


**Event-related potentials.** (**A**) Event-Related Potential (ERP) in the auditory condition for offset trials (dark blue) and constant trials (light blue) at electrode Cz. (**B**) ERP in the painful heat condition for offset trials (dark red) and constant trials (orange) at electrode Cz. The ERP was low-pass filtered at 7 Hz for visualization purposes.

**S12 Fig. Low-pass & high-pass filtered event-related potential (ERP) for auditory and thermal stimulation**

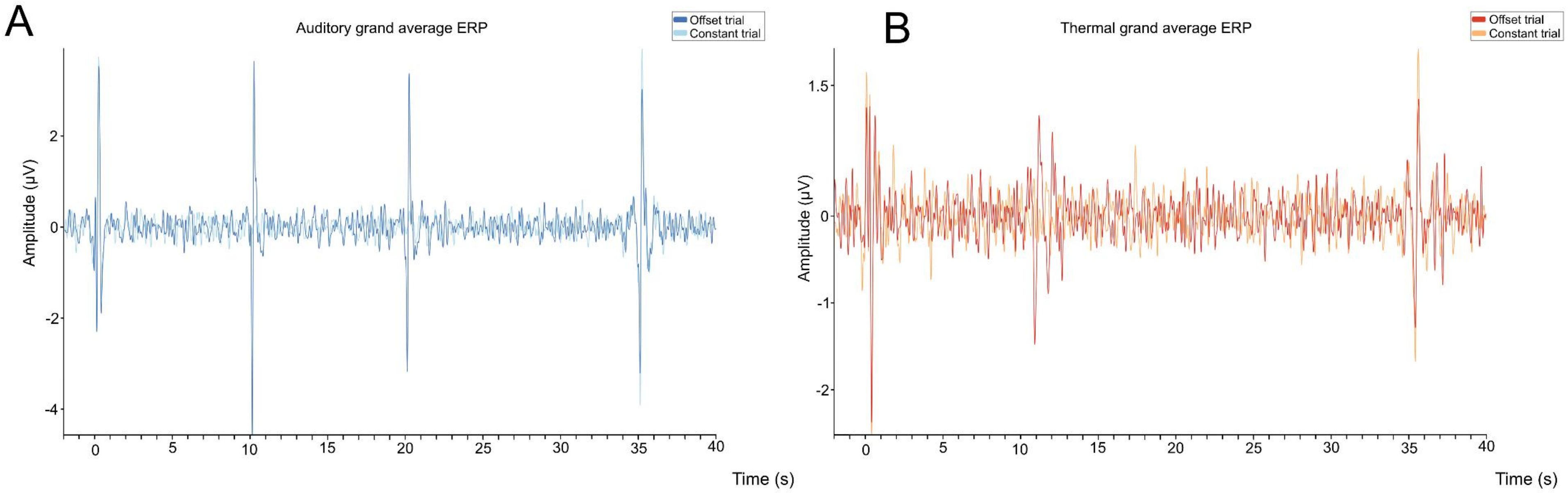


**Event-related Potentials.** (**A**) Event related Potentials (ERPs) in the auditory condition for offset trials (dark blue) and constant trials (light blue) at electrode Cz. (**B**) ERPs in the painful heat condition for offset trials (dark red) and constant trials (orange) at electrode Cz. The ERP was low-pass (7 Hz) and high-pass (1 Hz) filtered for visualization purposes.
